# High doses of fine biochar in sandy subsoils increase water retention, but also cause first-year yield depressions for drought-stressed barley

**DOI:** 10.1101/2025.10.09.681326

**Authors:** E. W. Bruun, C. T. Petersen, P. Iturbe-Espinoza, A. Winding, D. Müller-Stöver

## Abstract

**Context:** Coarse sandy subsoils often suffer from low water retention and rooting depth, limiting crop yields, especially under drought conditions.

**Aims:** To determine whether high doses of fine-grained biochar (150 or 300 Mg/ha) incorporated into sandy subsoil could improve water retention, nutrient uptake, root development and yield of spring barley under drought conditions.

**Methods:** A two-year mesocosm study (2022-2023) using soil columns containing subsoil layers of ground biochar (<80 µm, at 1%, 2% and 4%) or biochar pellets (2%) compared to unamended controls in two sandy soils.

**Key results:** Ground biochar increased available water capacity while intact pellets showed no measurable effect. First-year (2022) biochar applications negatively affected plant growth and nitrogen (N) uptake, likely due to N immobilization. Second-year (2023) results showed neutral to positive yield effects following winter irrigation. Biochar addition did not improve phosphorus uptake, despite the creation of wetter subsoils enriched with P. Root density decreased with increasing biochar concentration especially in one soil type, possibly due to increased potassium levels. Rooting depth was unaffected and greater than under field conditions.

**Conclusions:** Biochar incorporation into sandy subsoils can improve water retention, but may cause temporary N immobilization, affecting first-year crop performance. Overall benefits may emerge in subsequent growing seasons.

**Implications:** Long-term field trials are needed to assess nutrient management strategies, including additional N-application the first year. Incorporating biochar in autumn may help avoid N immobilization before spring sowing. Field-scale methods for applying fine-grained biochar to sandy subsoil must be developed.

**Highlights:**

1. Fine-grained biochar can increase the *in-situ* water retention in coarse sandy subsoils
2. In the first season (2022), biochar reduced barley growth and yield, likely due to initial soil nitrogen immobilization
3. In the second season (2023), after intensive winter irrigation, biochar effects became neutral or positive
4. Root density decreased with biochar use, possibly due to high potassium levels creating unfavourable subsoil conditions

## INTRODUCTION

Biochar, a carbon-rich recalcitrant material derived from the pyrolysis of organic matter, has received considerable attention for its ability to sequester carbon and improve soil fertility (Lehmann & Joseph, 2015). As a promising tool for climate change mitigation, biochar is expected to play an increasingly important role in the near future. Industrial scale pyrolysis facilities already exist or are under development in several countries. In Denmark, biochar is expected to play a key role in the green transformation of the agricultural sector (Skatteministeriet 2024), with the aim of producing thousands of tons of biochar.

However, the impact of biochar on soil fertility varies significantly across different soil types and climatic conditions. This variability highlights the need for targeted application to ensure a positive outcome. While results on clay soils are more inconsistent, research has shown that biochar can significantly improve the properties of sandy soils (Joseph et al. 2021). Agricultural land with coarse sandy soils faces many challenges, mainly due to low water retention, shallow root growth, and poor nutrient availability in the subsoil. These problems are widespread throughout the world, including in Denmark, where a quarter (24.5%) of the total land area consists of coarse sandy soils. As future climates are expected to have more frequent and severe droughts and periods of excessive rainfall, management of coarse sandy soils may become even more challenging, with increased irrigation requirements and risk of nutrient leaching. Historically, efforts have been made to improve sandy subsoils (Andersen 1985 and references therein), such as the addition of large amounts of sphagnum (Øvig 1979) or straw (Larsen & Øvig 1979) and mechanical subsoil loosening (Munkholm, Schjønning, and Sørensen 2003; Munkholm et al. 2005). In Australia, significant national research initiatives have investigated practical solutions to reduce subsoil constraints (Gill et al. 2012 and references therein). However, the beneficial effects of these methods typically diminish or disappear after a few years, so more permanent solutions are needed to improve sandy subsoils.

More recently, the incorporation of fine-grained biochar into the subsoil below the plough layer has been proposed (Bruun et al. 2014, 2021). This method aims to increase crop yields by directly improving subsoil conditions in the long term. Incorporation of fine-grained biochar into the coarse sandy subsoil can strongly affect water retention and hydraulic conductivity, as small biochar particles fill into large drainable soil pores, converting these pores into smaller water-retaining pores (Bruun et al. 2023, Petersen et al. 2016). This can also potentially lead to an increase in the retention and utilization of nutrients, thereby reducing leaching. Moreover, it has been shown that the application of fine biochar to the subsoil at doses of 1-2% can stimulate root proliferation, although at 4%, biochar had detrimental effects (Bruun et al. 2014).

Unlike organic materials such as manure or straw, which need to be replenished annually or more frequently when used as an amendment, the recalcitrant nature of biochar offers great potential for high carbon sequestration and long-lasting effects after just one large application. From a practical point of view, the farmer would benefit from applying biochar once rather than every year. From a carbon sequestration point of view, it would be beneficial if the application were as large as possible to sequester more carbon and thereby mitigate climate change the most. However, the potential effects (beneficial or detrimental) on crops following subsoil incorporation of a large amount of biochar into coarse sandy soil are largely unknown. Some studies have shown negative initial crop yields after biochar applications and attributed this to nutrient deficiencies caused by the short-term immobilization of nitrogen induced by biochar, as biochar typically contains (small) amounts of labile carbon (Bruun et al. 2012, Hansen et al. 2015). Excessive increases in soil pH and salinity after the application of large amounts of biochar may also potentially inhibit plant growth (Brtnicky et al. 2021). However, these effects are expected to be temporary, especially in sandy soils, as water-soluble salts released from the biochar are leached out of the soil profile. Leaching could potentially be problematic in terms of phosphorus loss, which may result in the eutrophication of waterways. However, biochars, particularly those produced from P-poor feedstocks and at higher pyrolysis temperatures, have been found to contain only small amounts of readily soluble P. Moreover, the release of P from these materials is highly dependent on several factors, such as pH and calcium (Ca) present in the soil solution or biochar (Buss et al. 2018, Robinson et al. 2018). However, a slow release of P in the subsoil could be beneficial for plant P uptake during periods when the topsoil is dry.

The objective of this study was therefore to assess whether spring barley grown on coarse sandy soil and under drought stress could benefit from the incorporation of high doses of fine-grained biochar (or biochar pellets) into the sandy subsoil. The primary aim was to evaluate plant responses to water and nutrient availability and to verify the (positive) effects on subsoil hydrology observed in laboratory and semi-field experiments (Bruun et al. 2014, 2021, 2023).

Overall, we hypothesized that fine-grained biochar would improve the use of water and nutrient resources and thereby increase the yield during periods of drought. To investigate this, we assessed several parameters including crop growth, yields, nutrient content in soil, water and nutrient uptake by plants, and root distribution, as well as physicochemical properties of the soil.

We specifically tested the following hypotheses:

i. Incorporation of high doses of finely ground biochar pellets (equivalent to 150 to 300 tonnes/ha) into coarse sandy subsoil will increase the available water capacity (AWC) and the effects will be greater than with equivalent amounts of intact pellets
ii. In the initial period (months) following the application of high doses of biochar, there will be adverse effects on plant growth due to changes in soil salinity and/or nitrogen immobilization
iii. Accelerated percolation equivalent to three years of net precipitation can eliminate or reduce any negative effects of high doses of biochar

## MATERIALS AND METHODS

### Soil and biochar

The experiment used topsoil (0-30 cm) and subsoil (30-100 cm) obtained from agricultural fields at two different locations: St. Jyndevad Experimental Station (54°53’N, 9°07’E) and near Billund (55°75’N, 9°02’E), Denmark. These sites, approximately 100 km apart, were chosen to provide variations of the same soil type. Both soils are characterized as “Coarse Sandy Soil” (JB1) according to the Danish soil classification system and as Spodosols (Typic Haplohumod) according to the USDA Soil Taxonomy system (Møberg et al. 1988). The Billund soil contains a slightly higher proportion of clay, silt, and fine sand compared to the Jyndevad soil. The textural composition and soil organic matter of both soils are detailed in Table 1 and Bruun et al. 2023.

**Table 1.** The textural composition (0-2000 µm ex. stones), percentage of stones and soil organic matter of the two subsoils used in the experiment (Bruun et al. 2023).

| Soil texture | Fraction | Jyndevad soil | Billund soil |
| --- | --- | --- | --- |
|  | µm | % | % |
| Clay | 2 | 1.5 (0.2) | 1.9 (0.8) |
| Silt | 2-20 | 1.4 (0.1) | 2.2 (0.4) |
| Fine sand | 20-50 | 0.3 (0.4) | 0.9 (0.7) |
|  | 50-100 | 1.7 (0.3) | 4.2 (0.2) |
|  | 100-200 | 8.7 (0.7) | 11.2 (0.4) |
| Coarse sand | 200-250 | 7.4 (0.3) | 5.9 (0.5) |
|  | 250-400 | 24.9 (0.4) | 24.3 (0.4) |
|  | 400-500 | 11.3 (0.7) | 12.9 (2.1) |
|  | 500-1000 | 35.8 (2.6) | 26.1 (1.9) |
|  | 1000-2000 | 7.0 (0.6) | 10.6 (0.3) |
| Stones | >2000 | 4.2 (0.2) | 8.7 (1.3) |
| SOM | - | 1.0 (0.1) | 0.7 (0.3) |

The biochar was produced by slow pyrolysis in a 200 kW pilot-scale plant (Stiesdal, Denmark) at approximately 550 °C and with a residence time of 30 minutes using straw pellets as feedstock. Some of the biochar pellets were afterwards finely ground in a hammer mill optimized for fine grinding (Euromill, Denmark) and equipped with an 80 µm screen. The particle size distribution (on a volume basis) of the fine biochar and soil was obtained with a Mastersizer3000 (Malvern Panalytical, United Kingdom) (Figure s-1), and the elemental composition of the biochar was determined by an elemental analyser (EA3000, Eurovector, Italy). The biochar contained 73.1 % carbon (C), 2.3 % hydrogen (H), 6.2 % oxygen (O), 1.1 % nitrogen (N) and 0.2 % P. The content of nutrients added per kg of subsoil can be seen in Table s-1. The content of total PAHs (total 16 EPA-PAH, analysed by Eurofins, Germany) was 3.7 mg/kg dw (Table s-2), which is below the threshold (6.0 mg/kg dw) applied in the EBC guidelines for agricultural systems (EBC (2012-2023)).

### Experimental setup

The two-year biochar experiment was conducted at the University of Copenhagen’s experimental farm Højbakkegaard in Taastrup (55°67′N, 12°31′E) with an experimental setup modified after Bruun et al. (2014) and (2021). The experiment consisted of nine treatments (Table 2). One group with irrigation was conducted in Jyndevad soil (J-), and two non-irrigated ‘drought’ groups were conducted in either Jyndevad or Billund soil (B-).

**Table 2.** List of the nine treatments used in the column experiment. Fine biochar had an average median diameter of 33 µm. J = Jyndevad soil, B = Billund soil.

| Treatment name | Soil (J-, B-) | Biochar concentration | Irrigation (-IR) | Biochar particle size (avg.) |
| --- | --- | --- | --- | --- |
| J-0%-IR | J | 0% | + | - |
| J-2%-IR | J | 2% | + | 33 µm |
| J-0% | J | 0% | - | - |
| J-1% | J | 1% | - | 33 µm |
| J-2% | J | 2% | - | 33 µm |
| J-4% | J | 4% | - | 33 µm |
| J-2%-PEL | J | 2% | - | 8x5 mm (pellets) |
| B-0% | B | 0% | - | - |
| B-2% | B | 2% | - | 33 µm |

The irrigated treatments (-IR) included: control (J-0%-IR) and amendment with 2% fine biochar (J-2%-IR). The non-irrigated treatments in the Jyndevad soil included: control (J-0%), amendment with 1%, 2% and 4% fine biochar (J-1%, J-2%, J-4%, respectively), and amendment with 2% biochar pellets (J-2%-PEL). Finally, two non-irrigated treatments were carried out in the Billund soil: A control (B-0%), and amendment with 2% fine biochar (B-2%). The application of 4% biochar to the subsoil (25-75 cm layer) corresponds to 300 tons of biochar per hectare.

Thirty-two soil columns were prepared in PVC cylinders (internal diameter 30 cm, height 150 cm) (Figure 1). To minimize preferential root growth along the otherwise smooth column walls, the insides of the cylinders were coated with subsoil material using insoluble wallpaper glue, following the method described by Bruun et al. (2014 and 2021). Each column was uniformly packed with soil by adding air-dried soil volumes of 15-20 litres each, followed by thorough tapping on the sides to allow the soil to settle. Initially, untreated subsoil was added to each column to form a bottom layer of approx. 70 cm (80–150 cm below the upper rim). This layer was added to reduce the impact on the water content from the lower boundary, where a free water surface is formed whenever drainage occurs. Subsequently, a 50 cm layer of subsoil with or without biochar amendment was added (approximately 30–80 cm below the upper rim), followed by a 25 cm layer of the original topsoil (approximately 5-30 cm below the rim). The biochar-amended subsoil was carefully mixed using a pile mixing procedure before being added to the columns. An access pipe for water monitoring to a maximum depth of 100 cm was installed centrally in each column in late April 2022. The soil columns were placed on a tiled surface in an outdoor area that received sunlight but was sheltered from the wind. They were arranged in a randomized block design with four blocks to account for potential small variations in sunlight and wind exposure. Weather data were obtained from the university climate station located less than 1 km away.

**Figure 1.**
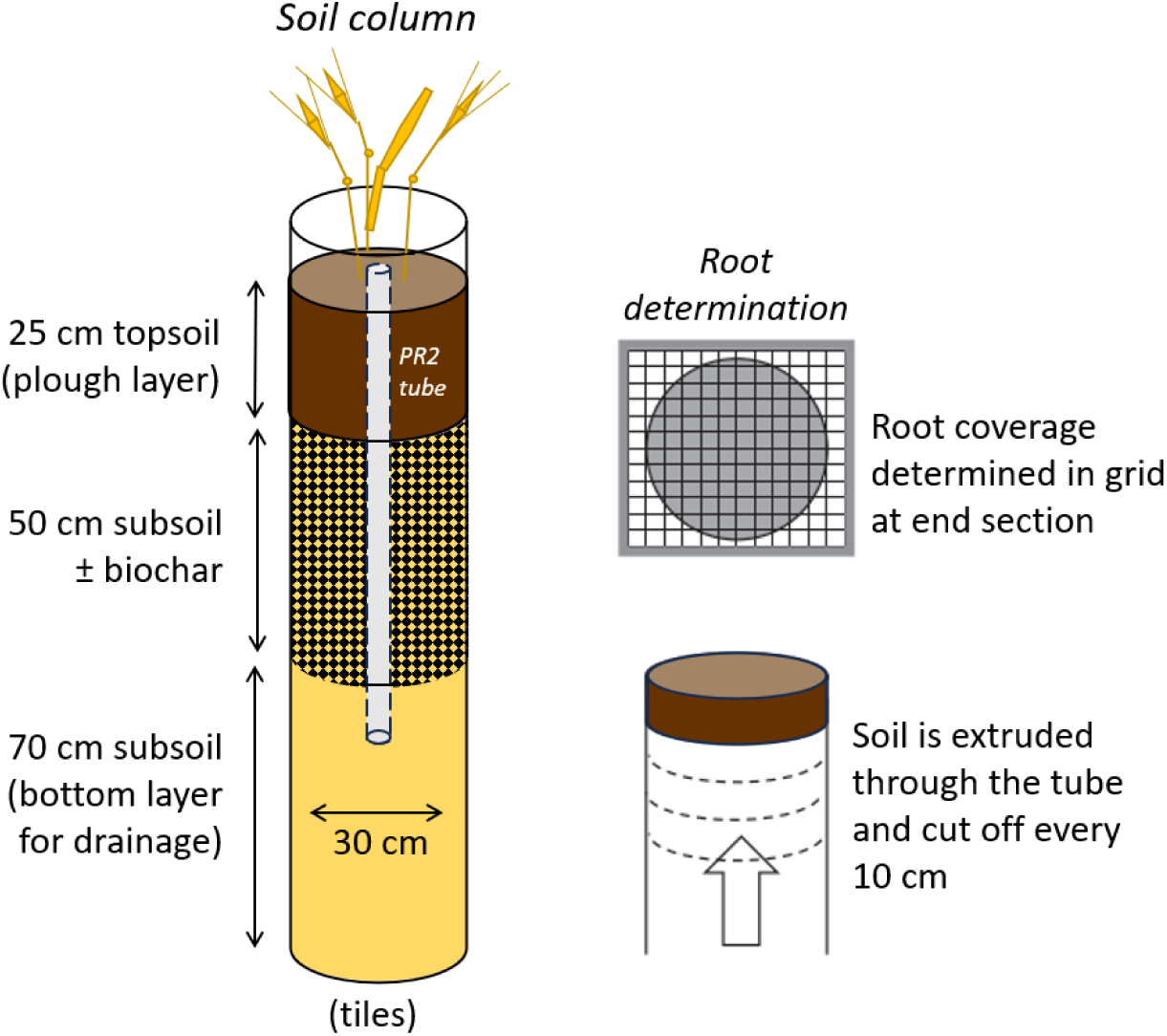
Left: Schematic diagram of one soil column with spring barley. The PR2 tube for water measurements is indicated. Right: methodology for determination of roots in the soil profile after crop harvest. Modified after Bruun et al. 2014.

### Crops, crop management, irrigation, and water measurements

The crop was spring barley (*Hordeum vulgare* L.) in both growing seasons, sown on April 19, 2022, and March 23, 2023. The number of plants per column was reduced to 20 (equivalent to 280 m^−2^) during the early tillering growth stage. A rain shelter roof was installed over the soil columns on June 1 in both years to ensure drought in the non-irrigated treatments (Figure s-2). In 2022, the roof was removed on June 30, while in 2023 the roof was left in place until harvest. Immediately after sowing, mineral NPK fertilizer (6.7 g 21– 3–10 per column, equivalent to 200 kg N/ha) was applied to the soil surface in both growing seasons. The crop was harvested on August 10, 2022, and July 13, 2023, respectively, by cutting the straw approximately three cm above the soil surface. Dry matter yield was determined separately for straw and grains after oven drying at 80 °C. Moreover, the height of the longest straw was measured in all soil columns during each growing season.

Irrigation was applied once or twice a week to avoid water deficits of more than approximately 35 mm in the treatments J-0%-IR and J-2%-IR. The amount of irrigation water applied was based on measuring the difference between the actual water content in the 0-105 cm soil layer and the estimated field capacity.

During the winter of 2022-2023, a thorough irrigation regime was applied to all soil columns to mitigate potentially detrimental short-term effects (e.g. salt stress) following the large addition of biochar. The columns received 1494 mm of water (precipitation plus irrigation) during the period November 1, 2022, to April 1, 2023, resulting in a percolation approximately equivalent to three years under natural field conditions at the Jyndevad site (Henriksen and Sonnenberg, 2003). The soil was left bare between the harvest and the following sowing.

Soil water content measurements using the PR2 profile probe (Delta-T Devices Ltd, Cambridge, UK) were carried out systematically in all columns during the growing seasons from May 2022 to August 2023. Measurements were taken at depths of approximately 10, 20, 30, 40, 50, 60, 90, and 100 cm, representing the respective soil layers: 0–15, 15–25, 25–35, 35–45, 45–55, 55–75, 75–95, and 95– 105 cm, following the methodology described in Bruun et al. (2021). At each depth, measurements were conducted in triplicate by turning the PR2 unit 120 degrees between each measurement. In addition to the PR2 water content measurements, three Teros 31 tensiometers (METER Group, Munich, Germany) were installed (from the column sides) in a non-irrigated control column (J-0%) and a column with 4% biochar amendment (J-4%) at 40, 60 and 90 cm depth to provide continuous measurements of matric potential.

The accumulated actual evapotranspiration (Ea) was estimated as described in Bruun et al. (2021), using the formula:

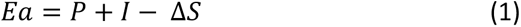

where *P* and *I* represent the accumulated precipitation and irrigation over a certain period, respectively. *ΔS* denotes the change in soil water content (from 0 to 105 cm depth) within the same period. The change in soil water content, *ΔS* was calculated as:

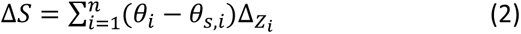

where θ_s,i_ and θ_i_ represent the measured volumetric water content in soil layer *i* at the start and end of the considered period, respectively. Δz_i_ is is the thickness of layer *i* and *n* is the number of layers (n = 8).

### Root inspection procedure, soil moisture, and bulk density determinations

Root inspection and further soil analyses were carried out in August 2023 on selected treatments (J-0%, J-2%, J-4%, B-0%, B-2%) at 40, 50, 60, 70, and 90 cm depth after the final harvest and thorough irrigation to determine field capacity with the methodology used by Bruun et al. (2014). A lifting jack system was employed to extrude the soil from the columns, while a thin steel plate facilitated the sequential slicing of the soil into 10 cm cylindrical segments (Figure 1). This procedure allowed the assessment of root distribution, bulk density, and soil moisture content at different depths in the soil profile. Soil moisture and bulk density determinations were based on undisturbed soil samples (2 x 100 cm^3^) in each slice. Root distribution was assessed using the grid-based method used by Bruun et al. (2014) and Munkholm et al. (2008). Briefly, the exposed and freshly cut end of the soil column was carefully brushed to remove excess soil, and a grid net with square grid cells measuring 2.5 by 2.5 cm (144 cells in total) was placed against the surface. All grid cells were visually inspected (under strong lighting) by the same person to determine the presence or absence of roots (yes or no). Subsequently, the root coverage for each depth examined was calculated by dividing the area of grid cells containing roots by the total cross-sectional area of the soil. Grid cells crossing the column rim, i.e. where the PVC column wall was visible, were assigned a weighting factor proportional to the soil area.

Bulk density (ρ_b_) and gravimetric soil moisture content (w) were determined by weighing before and after oven drying at 105°C. The volumetric water content was calculated using the formula:

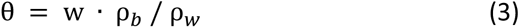

where ρ_w_ represents the density of water.

### Soil and crop analyses

Soil samples were collected from a depth of 50 cm in all treatments in 2022 (June and September) and 2023 (May and June) for analysis of exchangeable K, plant-available P, mineral N (nitrate + ammonium), pH, and electrical conductivity (EC). Exchangeable potassium in air-dried soil samples was extracted with 1 M ammonium acetate (pH 7) and analysed by ICP-OES (Agilent 5100; Agilent Technologies, Santa Clara, USA). For Olsen P extraction, soil samples were mixed with a 0.5 M sodium bicarbonate solution (pH 8.5), shaken for 30 minutes, filtered, and treated with sulfuric acid before analysis by Flow Injection Analysis (FIA) (FIAstar 5000 Analyzer, Foss, Sweden). Nitrate and ammonium were extracted with 1 M KCl (1:4 soil:KCl suspension), shaken for 60 minutes, filtered, and analysed by FIA. pH and EC were measured in a 1:5 suspension (soil:Milli-Q water), shaken for 60 minutes. The samples was filtered prior to EC measurements.

Ground straw and grain samples were analysed for their elemental composition by ICP-OES (Agilent 5100; Agilent Technologies) after microwave digestion in nitric acid and hydrogen peroxide and for their total N content on an elemental analyzer (Vario macro cube, Elementar Analysensysteme GmbH, Germany).

For the analysis of extracellular enzymatic α-glucosidase activity, soil samples were sieved, mixed with MQ water, and centrifuged according to the procedure described in Supplementary Materials, as described in Iturbe-Espinoza et al. (under review). Enzyme activity was expressed as nmMUF g⁻¹ dry soil h⁻¹.

### Statistical analysis

Levene’s test was used to test for homogeneity of variance. All data were analysed by one-way analysis of variance (ANOVA) using SPSS software (29.0.10). Data for biomass and straw height were analysed by one-way ANOVA with the four blocks as random factor. Tukey’s adjusted differences of least square means were used for multiple pairwise comparisons. In all statistical tests, a *p*-value of 0.05 or less was considered significant. Data from the two treatments with Billund soil and the two irrigated treatments were analysed using Fisher’s protected *t*-test.

## RESULTS

### Weather conditions and soil water balances

The two growing seasons differed in terms of weather conditions (Figure 2 A, B). The reference evapotranspiration (Penman, 1956) accumulated from May 1 to July 13 was 234 mm in 2022 but 273 mm in 2023, and the non-irrigated treatments received less water during this period in 2023 (6 mm in 2023 compared with 76 mm in 2022). Evapotranspiration, i.e. water extracted from the 0-105 cm layer, was similar between the different non-irrigated treatments, except for a (non-significant) tendency in 2023 for J-4% to have higher values than treatments with lower biochar concentrations.

**Figure 2.**
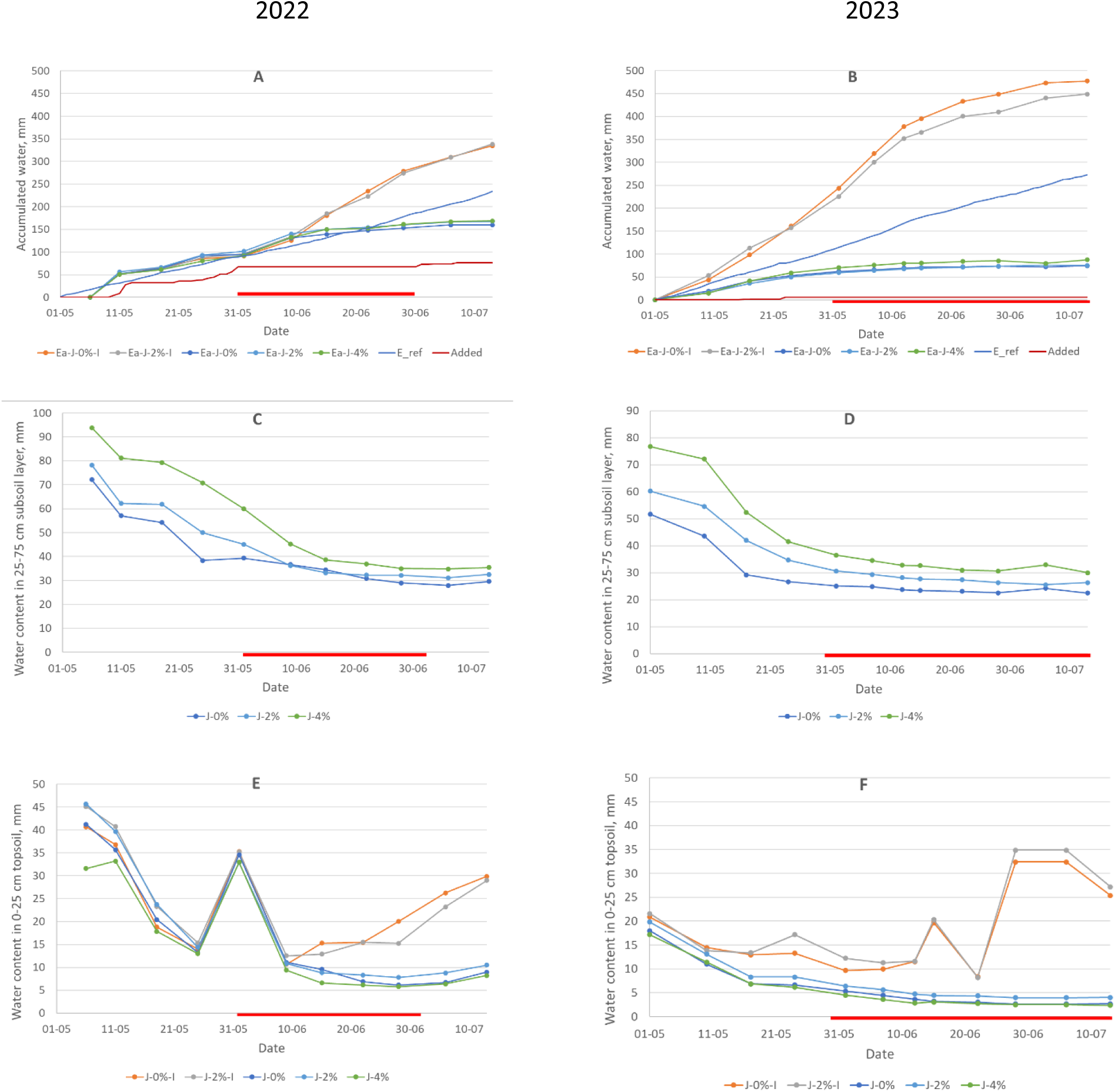
Water balances in selected treatments in 2022 (left) and 2023 (right). A and B: calculated actual evapotranspiration Ea (profile method, 0-105 cm depth) for the treatments J-0%, J-0%-I, J-2%-I, J-2%, and J-4% (accumulated values from May 6 in 2022 and May 1 in 2023). Reference evapotranspiration E_ref and added water (precipitation and irrigation (14 mm in 2022) given to all treatments) accumulated from May 1. The red bars indicate periods with a rain shelter. C and D: water content in the 25-75 cm treated subsoil layer for the treatments J-0%, J-2%, and J-4%. E and F: water content in the 0-25 cm untreated topsoil layer for the treatments J-0%, J-0%-I, J-2%-I, J-2%, and J-4%. Mean values of the four replicates. J- = Jyndevad soil, B- = Billund soil, -IR = irrigated, -PEL = pellet biochar, -x% = percentages of fine grained or pellet biochar. B-0%, B-2%, J-1% and J-2%-PEL have been excluded for clarity.

Evapotranspiration was higher in the treatments with optimal irrigation (J-0%-I and J-2%-I) than in the non-irrigated treatments. It was also considerably higher than the reference evapotranspiration (potential evapotranspiration from short-cut grass), likely due to oasis effects. The difference in actual evapotranspiration between the irrigated and non-irrigated treatments (as an expression of drought) was greater and occurred earlier in 2023 than in 2022.

The water content of the treated subsoil layer (25-75 cm depth) increased with biochar content (Figure 2 C, D). For the treatment J-4% at field capacity (approximately May 11, 2022, and May 1, 2023), it was 81 and 77 mm, corresponding to 16.2 and 15.4 vol%, respectively. For J-0% (the control) at field capacity, it was approximately 57 and 52 mm, corresponding to 11.4 and 10.4 vol%, respectively. Hence, adding 4 wt% of biochar increased the field capacity by approximately 24 and 25 mm over the two years, corresponding to 4.8 and 5.0 vol%. The water content decreased over time, reaching more or less stable final values at the end of June. These final values were lower in 2023 than in 2022, and they increased with increasing biochar content, indicating a slightly higher wilting point in soil rich in biochar. The application of 2 wt% of biochar in pellets (J-2%-PEL) had no significant effect on the field capacity of the two soils (not shown).

The water content of the topsoil (0-25 cm) reached very low values (7-8 mm) on May 17 in 2023, probably due to the very small amounts of water added in May before the establishment of the rain shelter, whereas similarly low values occurred one month later on June 15 in 2022. Hence, the non-irrigated crop experienced a much drier topsoil in the middle part of the growing season in 2023 than in 2022.

### Crop parameters

#### Crop development and yields

Spring barley yields showed considerable variability between years and between treatments (Figure 3 and Table 3). Across all treatments and in both years, the highest grain and straw yields were consistently and as expected observed in the two irrigated treatments (J-0%-IR and J-2%-IR). In 2022, the irrigated control yielded 10% more grain than the irrigated biochar treatment, whereas in 2023, grain yields were comparable.

**Figure 3.**
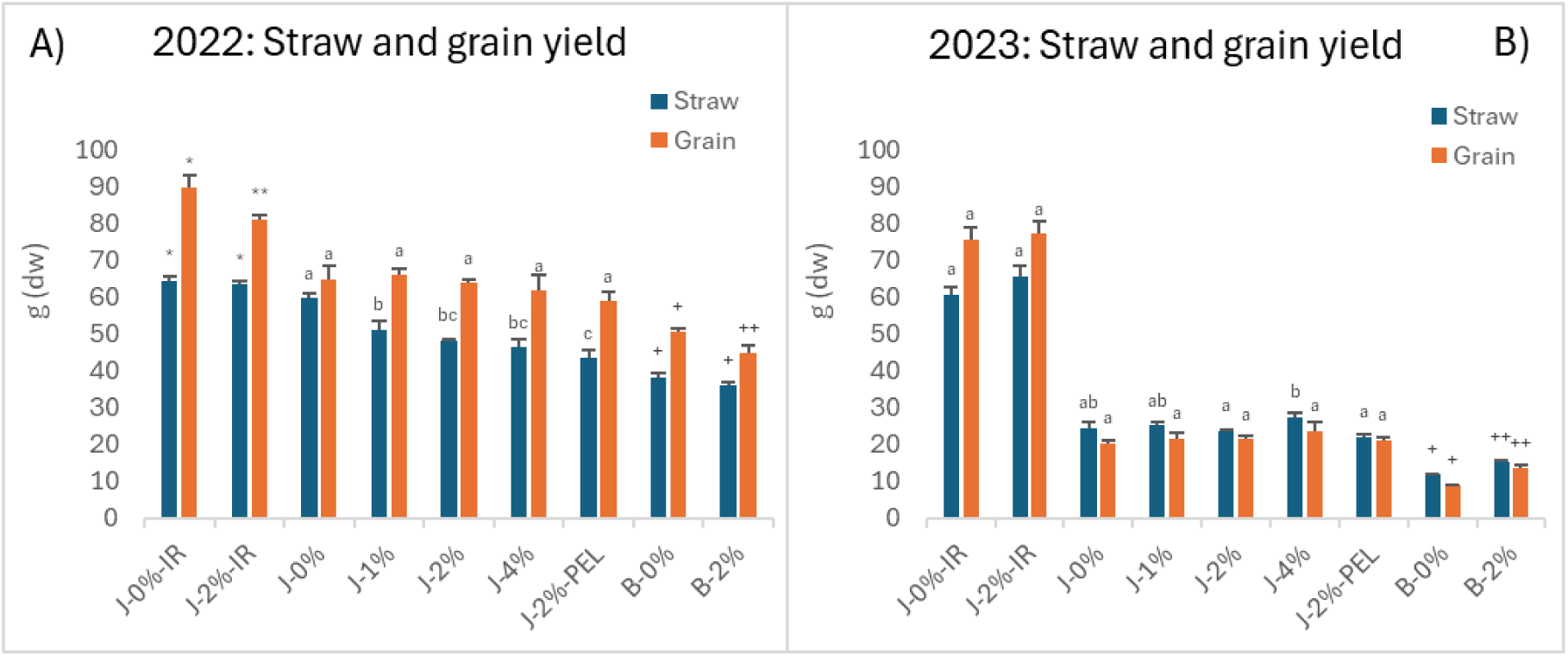
Average dry matter yield in straw and grain (g dry weight) in the nine treatments in A) 2022 and B) 2023. Mean ± standard error. Means with different letters or number of same symbols indicate significant differences at p = 0.05 within the same sampling date. Means were tested within the three groups: irrigated (-IR), non-irrigated in Jyndevad soil (J-), and Billund soil (B-). J- = Jyndevad soil, B- = Billund soil, -IR = irrigated, -PEL = pellet biochar, -x% = percentages of fine grained or pellet biochar.

**Table 3.**
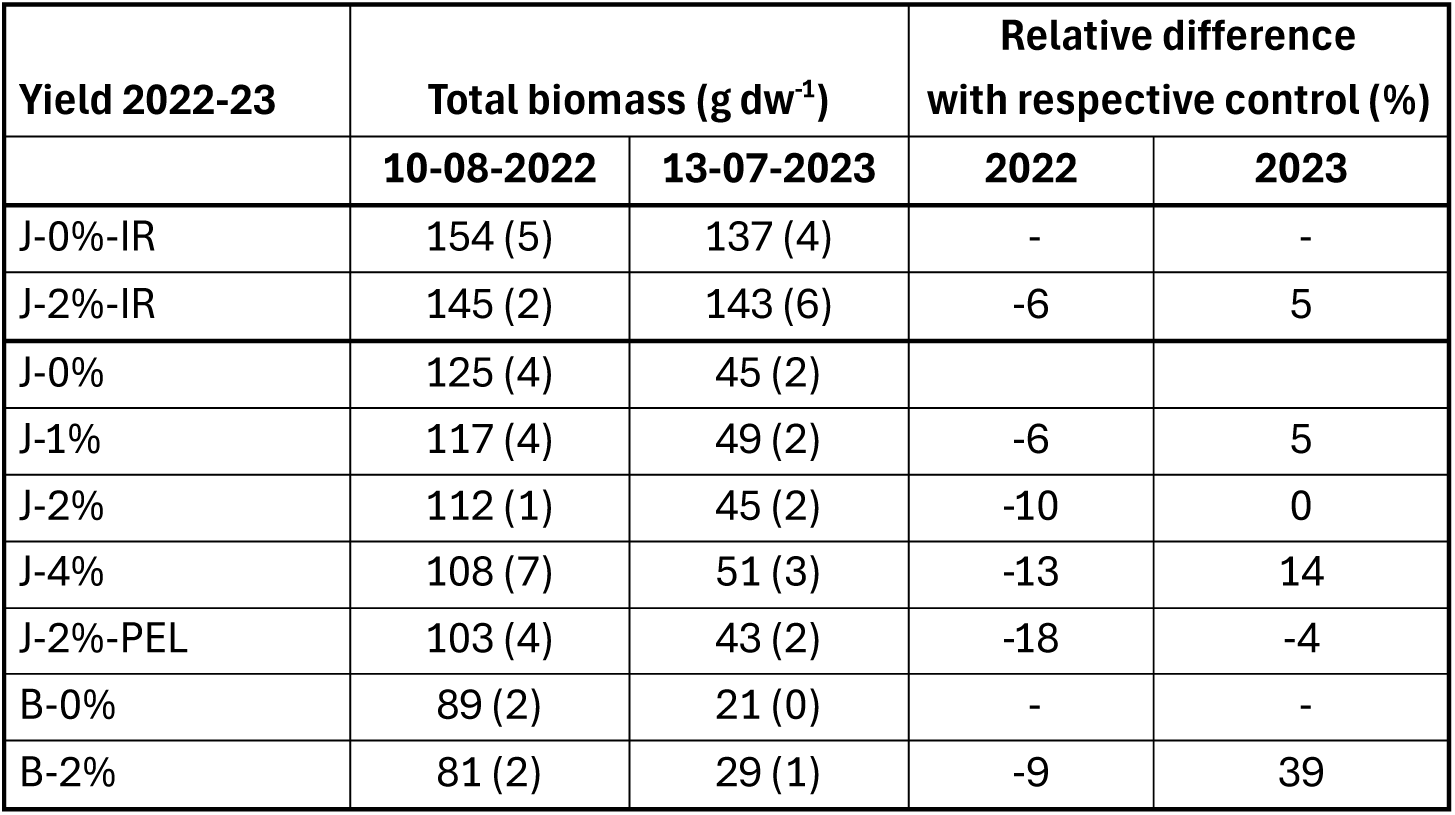
Total dry matter yield for the ten treatments in 2022 and 2023, and the relative difference within each year between the biochar treatments and their respective controls. Numbers in parenthesis are standard errors. J- = Jyndevad soil, B- = Billund soil, -IR = irrigated, -PEL = pellet biochar, -x% = percentages of fine grained or pellet biochar.

| Yield 2022-23 | Total biomass (g dw<sup>-1</sup>) |  | Relative difference with respective control (%) |  |
| --- | --- | --- | --- | --- |
|  | 10-08-2022 | 13-07-2023 | 2022 | 2023 |
| J-0%-IR | 154 (5) | 137 (4) | - | - |
| J-2%-IR | 145 (2) | 143 (6) | -6 | 5 |
| J-0% | 125 (4) | 45 (2) |  |  |
| J-1% | 117 (4) | 49 (2) | -6 | 5 |
| J-2% | 112 (1) | 45 (2) | -10 | 0 |
| J-4% | 108 (7) | 51 (3) | -13 | 14 |
| J-2%-PEL | 103 (4) | 43 (2) | -18 | -4 |
| B-0% | 89 (2) | 21 (0) | - | - |
| B-2% | 81 (2) | 29 (1) | -9 | 39 |

Yields in all non-irrigated treatments were notably higher in 2022 than in 2023, when crop development was significantly affected by the more severe drought. For instance, the biomass yields of the two non-irrigated controls were 64% (J-0%) and 77% (B-0%) lower compared to the previous year’s crop.

In 2022, there was a consistent trend across all treatments indicating decreasing biomass yields with increasing levels of biochar in both subsoils (Figure 3). Specifically, the straw yield in the J-0% treatment was significantly higher than that of all biochar drought treatments. This trend was further reflected in the stunted plant growth relative to the control observed in all biochar treatments in 2022. The measured height of the longest shoot showed a significant reduction in the biochar treatments compared to their respective controls (J-0%-IR, J-0%, B-0%), with a trend towards shorter “tallest shoots” as biochar concentrations increased (Figure s-3). Additionally, the stems in the biochar treatments showed a reddish colour, a phenomenon observed to a lesser extent in the controls (Figure s-4).

The results in the second year (2023) were different. In this year, the average biomass yields and shoot heights in all biochar treatments were similar to (and tended to be higher than) the respective controls. E.g. J-4% had a 12 % and 16 % higher straw and grain yields, respectively, than the Jyndevad drought control (J-0%). This was even more pronounced for the Billund soil, where B-2% had a significantly higher straw and grain yield than the control (B-0%) by 28 % and 53 %, respectively. However, compared to the irrigated treatments (J-0%-IR), the yields were still very low.

#### Straw and grain analyses

The mineral composition of both straw and grains showed variability within treatments and between years (Table s-3 and s-4). In 2022, a decrease in grain nitrogen (N) concentrations was observed in the biochar-amended treatments (J-4%, J-2%-PEL, and B-2%) compared to their respective drought controls (J-0% and B-0%), declining from 1.5% N in J-0% to 1.2% N in J-2%-PEL, and from 1.6% in B-0% to 1.2% N in B-2% (Figure 4AB). This decrease in grain N concentration coincided with lower biomass yields in these treatments compared to the drought controls (Figure 3). Thus, there was an overall greater total amount of N assimilated by the crops in J-0% and B-0% (Figure 4CD). In 2023, both straw and grains had increased N concentrations compared to the previous year. The irrigated treatments had markedly lower N concentrations compared to the drought treatments this year, but at the same time, these treatments had a much higher total N uptake per column as a result of the higher biomass yields. In contrast to 2022, no significant differences in N concentrations or N uptake were observed between the non-irrigated controls and the respective biochar treatments, except for the grain N concentration of J-2%-PEL which was again significantly lower than J-0%.

**Figure 4.**
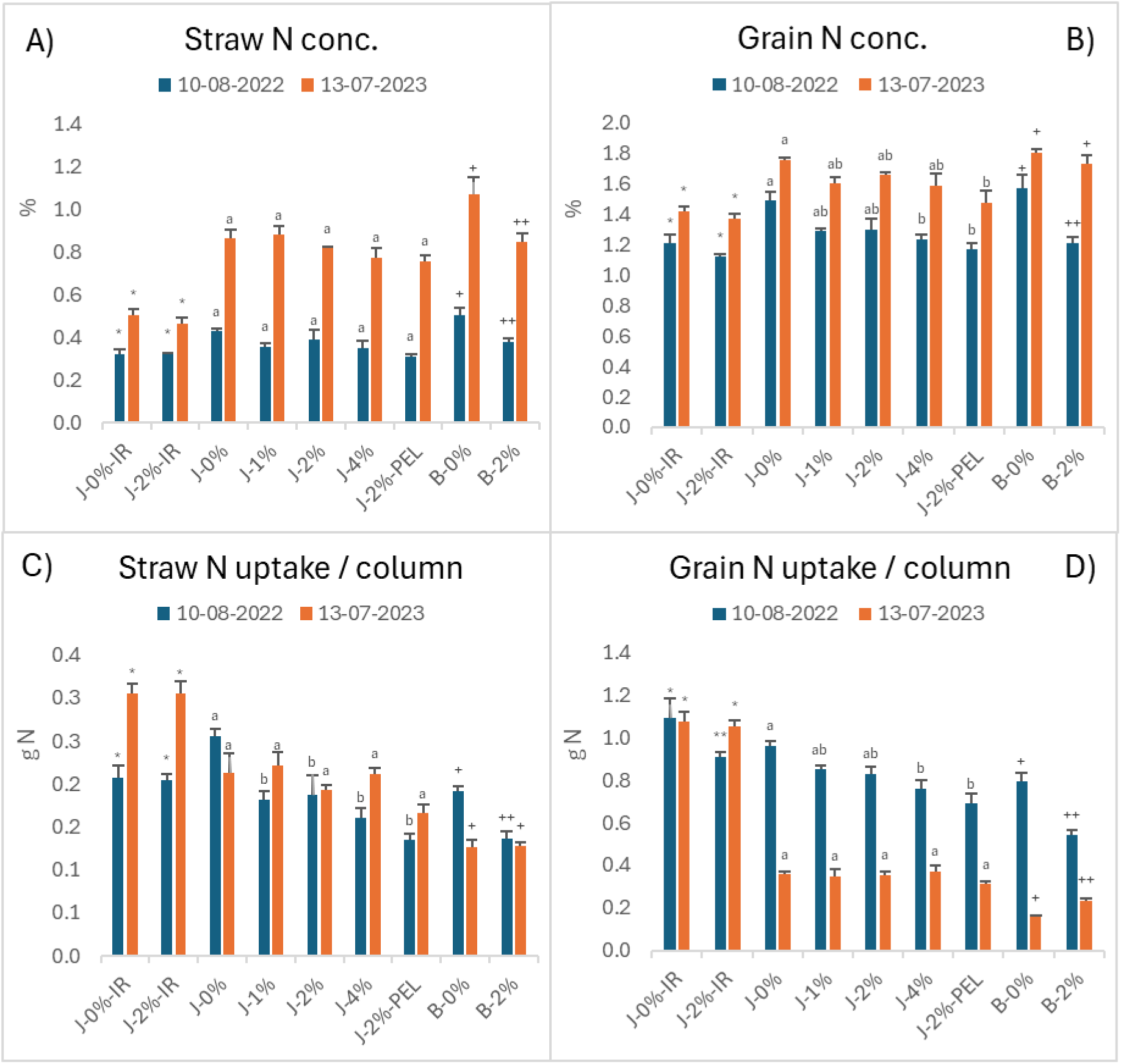
A) Concentration of nitrogen in straw (%). B) Concentration of nitrogen in grain (%). C) Total nitrogen uptake (g N) in straw pr. column. D) Total nitrogen uptake in grain pr. column (g N). Mean ± standard error. Means with different letters or number of same symbols indicate significant differences at p = 0.05 within the same sampling date. Means were tested within the three groups: irrigated (-IR), non-irrigated in Jyndevad soil (J-), and Billund soil (B-). For treatment abbreviations see figure 3.

Biochar had no clear effect on the phosphorus (P) concentration in straw and grain in both years, neither in the irrigated nor in the non-irrigated treatments. In 2023, the P concentration in the non-irrigated treatments was markedly decreased compared to the previous year and to the irrigated treatments (Figure 5).

**Figure 5.**
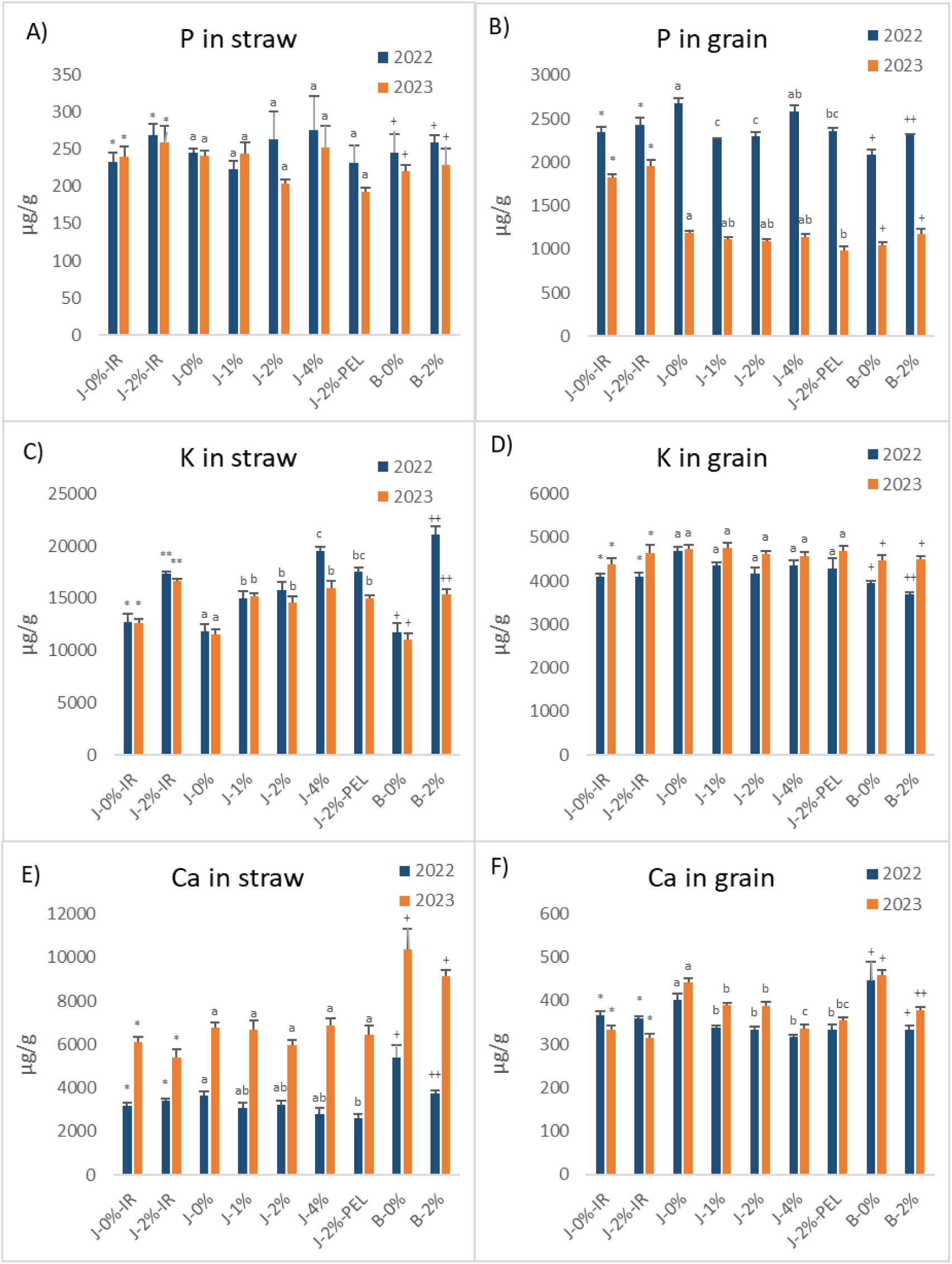
Average content (µg g^-1^) in straw and grain of phosphorus in A) and B), potassium in C) and D) and Calcium in E) and F). Straw and grains sampled after harvest 13. June 2022 and 10. August 2023. Mean ± standard error. Means with different letters or number of same symbols indicate significant differences at p = 0.05 within the same sampling date. Means were tested within the three groups: irrigated (-IR), non-irrigated in Jyndevad soil (J-), and Billund soil (B-). For treatment abbreviations see figure 3.

Biochar application increased potassium (K) concentrations in straw, with higher biochar levels corresponding to higher K concentrations. In 2023, K concentrations were lower, particularly in the non-irrigated treatments. In grain, K concentrations remained constant across treatments and years. Calcium (Ca) content varied between years, with higher concentrations measured in 2023 compared to 2022. In non-irrigated treatments, Ca concentrations in both straw and grain decreased with biochar application. In grain, Ca concentrations showed minimal interannual variation and were consistently higher in control soils under non-irrigated conditions.

#### Root density

In the treatments conducted in Jyndevad soil (J-X%), the root density expressed as degree of root coverage decreased noticeably with increasing biochar concentration (J-2%, J-4%) compared to the Jyndevad drought control (J-0%) (Figure 6). Furthermore, within each biochar treatment, root coverage decreased sharply with depth, e.g. from 51% at 40 cm depth to appr. 10% coverage in 90 cm depth in J-4%. J-0%, on the other hand, maintained a high constant root coverage of around 90% at 40, 50, 60 and 70 cm depth, and still had almost three times higher root coverage (28%) at 90 cm depth compared to all other treatments. In the Billund soil, the difference between the control and the biochar treatment was less clear, with B-2% tending to have a higher average root coverage at 40, 50 and 60 cm depth, while B-0% had the highest coverage at 70 and 90 cm depth.

**Figure 6.**
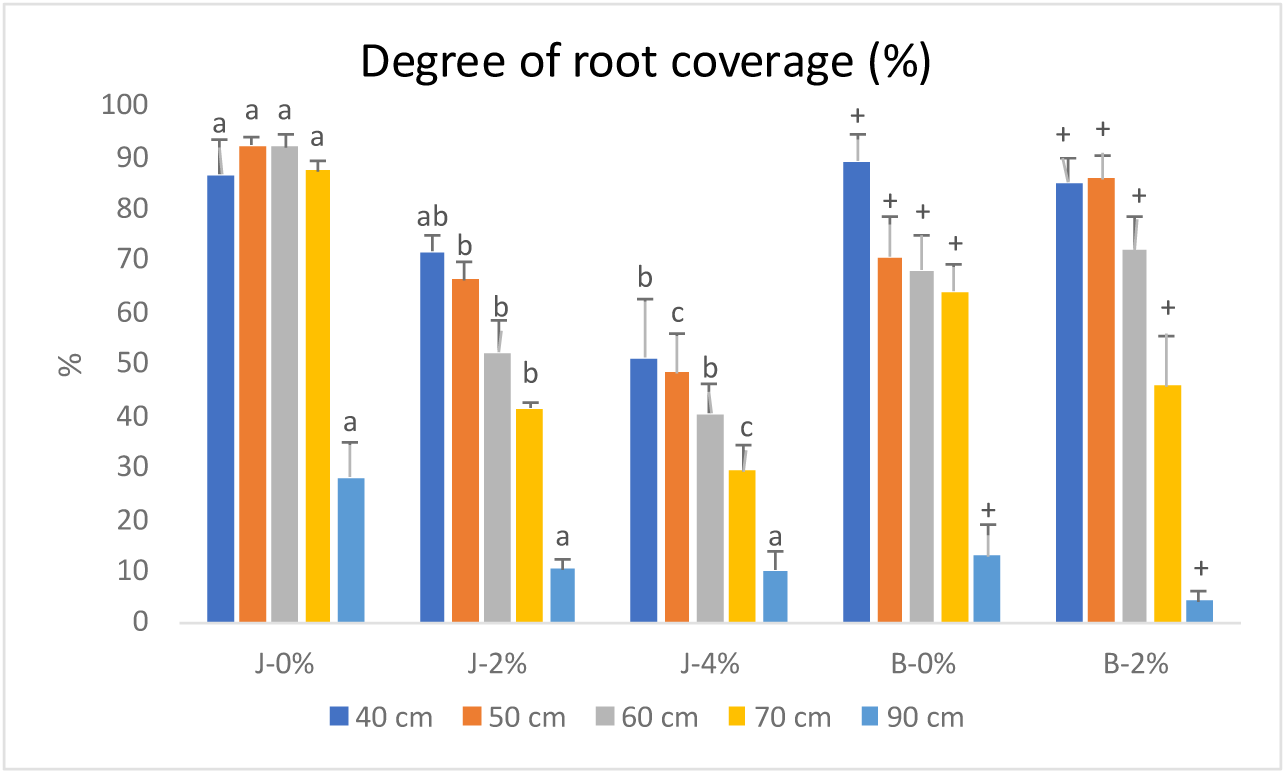
Degree of root coverage (%) of barley roots in coarse sandy subsoil amended with different amounts of straw-biochar measured at 40, 50, 60, 70, and 90 cm’s depth. Mean ± standard error. Means with different letters or number of same symbols indicate significant differences at p = 0.05 within the same sampling date. Means were tested within the two groups: Jyndevad soil (J-), and Billund soil (B-). For treatment abbreviations see figure 3.

In all treatments, we observed roots at the very bottom of the columns (at a depth of 150 cm), showing that the crops were able to extract water down to the artificial lower boundary. On July 13 in 2023, the water content was measured in all columns at a depth of approximately 130 cm (soil samples were taken 15 cm above the tiled ground surface in each column). The measured water content was very low, ranging from 2.2 wt% in J-0% to 5.3 wt% in B-2% (Figure s-5).

### Soil parameters

#### Electrical conductivity and pH

The electrical conductivity at 50 cm depth after the addition of biochar to the Jyndevad and Billund soils was only affected at the first sampling date (28-06-22), where the irrigated treatment (J-2%-IR), the Billund soil treatment (B-2%) and J-4% in the Jyndevad drought treatments had a significantly higher EC than their respective controls (Figure 7A). In the second growing season (2023) after the accelerated irrigation campaign in the winter 2022-23, the conductivity decreased in most cases. There was little change in pH over time, except at the start of 2023 (04-05-23). The pH increased moderately with increasing biochar concentration, reaching a pH of 7.3 in J-4% (Figure 7B).

**Figure 7.**
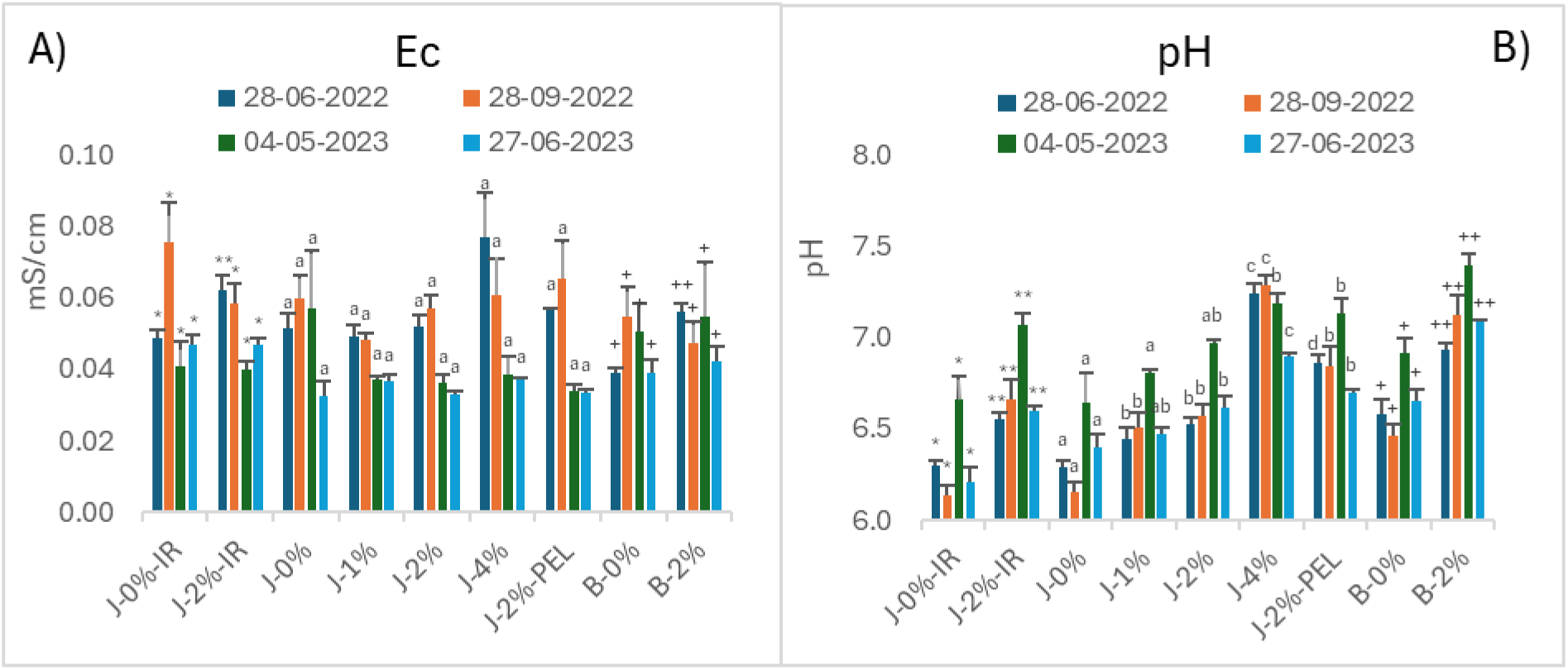
A) EC in a 1:5 (soil:water) suspension and B) pH of the nine treatments in 50 cm’s depth measured at four different times during the course of the experiment. Mean ± standard error. Means with different letters or number of same symbols indicate significant differences at p = 0.05 within the same sampling date. Means were tested within the three groups: irrigated (-IR), non-irrigated in Jyndevad soil (J-), and Billund soil (B-). For treatment abbreviations see figure 3.

#### Exchangeable Potassium (K)

The level of exchangeable K (15 to 29 mg/kg soil) in the control soils was not particularly high in any of the measurements conducted over the two years (Figure 8A). With increasing biochar concentrations, the K concentration increased up to a maximum of 319 mg kg^-1^ in J-4% at the first sampling date (28-06-2022). From 2022 to 2023 there was a large decrease in exchangeable K levels in all Jyndevad soil (J-X%) treatments, most pronounced in J-4%, likely due to the irrigation during the winter of 2022-23.

**Figure 8.**
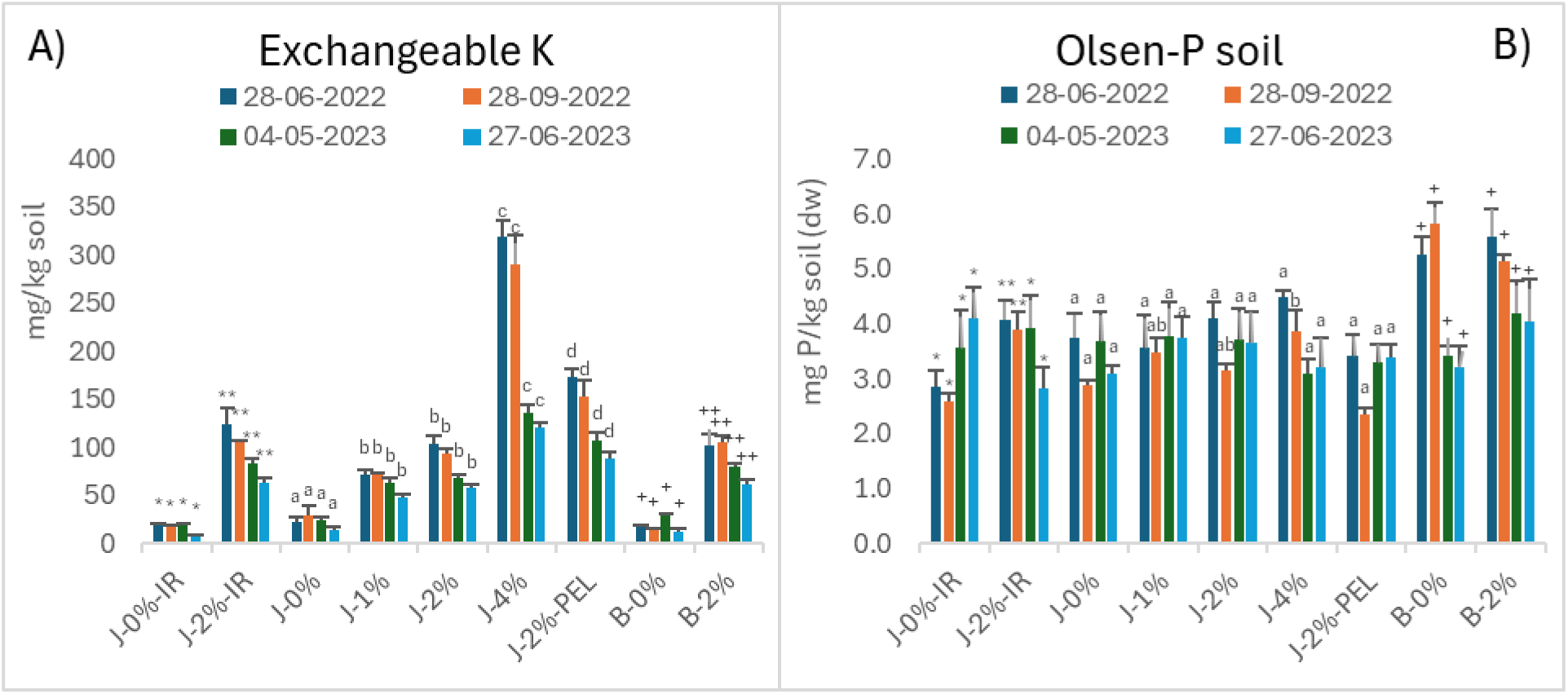
A) Exchangeable potassium (K) in a 1:5 (soil:water) suspension and B) Olsen-P of the nine treatments in 50 cm’s depth at four times during the experiment. Mean ± standard error. Means with different letters or number of same symbols indicate significant differences at p = 0.05 within the same sampling date. Means were tested within the three groups: irrigated (-IR), non-irrigated in Jyndevad soil (J-), and Billund soil (B-). For treatment abbreviations see figure 3.

#### Olsen-P

The Olsen-P content was overall low, and biochar did not significantly increase it in any of the years (Figure 8B). Contrary to the trends observed for exchangeable potassium and electrical conductivity, there was no significant decrease after winter irrigation in 2022-23, except in B-0%.

#### Microbial activity in soil (extracellular α-glucosidase)

The activity of α-glucosidase was significantly higher in the organic-rich surface soil (data not shown, see Iturbe-Espinoza et al. 2025 (in review)) compared with the lower depths of the soil column. In the subsoil at the end of the experiment (August 2023) the average α-glucosidase activity was 0.60, 0.62 and 97 nmMUF g^-1^ dry soil h^-1^ for the J-0%, J-2% and J-4%, respectively, while in the Billund soil, it was 0.42 and 0.51 nmMUF g^-1^ dry soil h^-1^ for the B-0% and B-2% respectively (Figure 9). Biochar amendment increased extracellular α-glucosidase activity in both subsoils, with significant differences between the higher concentration J-4% and either J-0% or J-2% in the Jyndevad subsoil.

**Figure 9.**
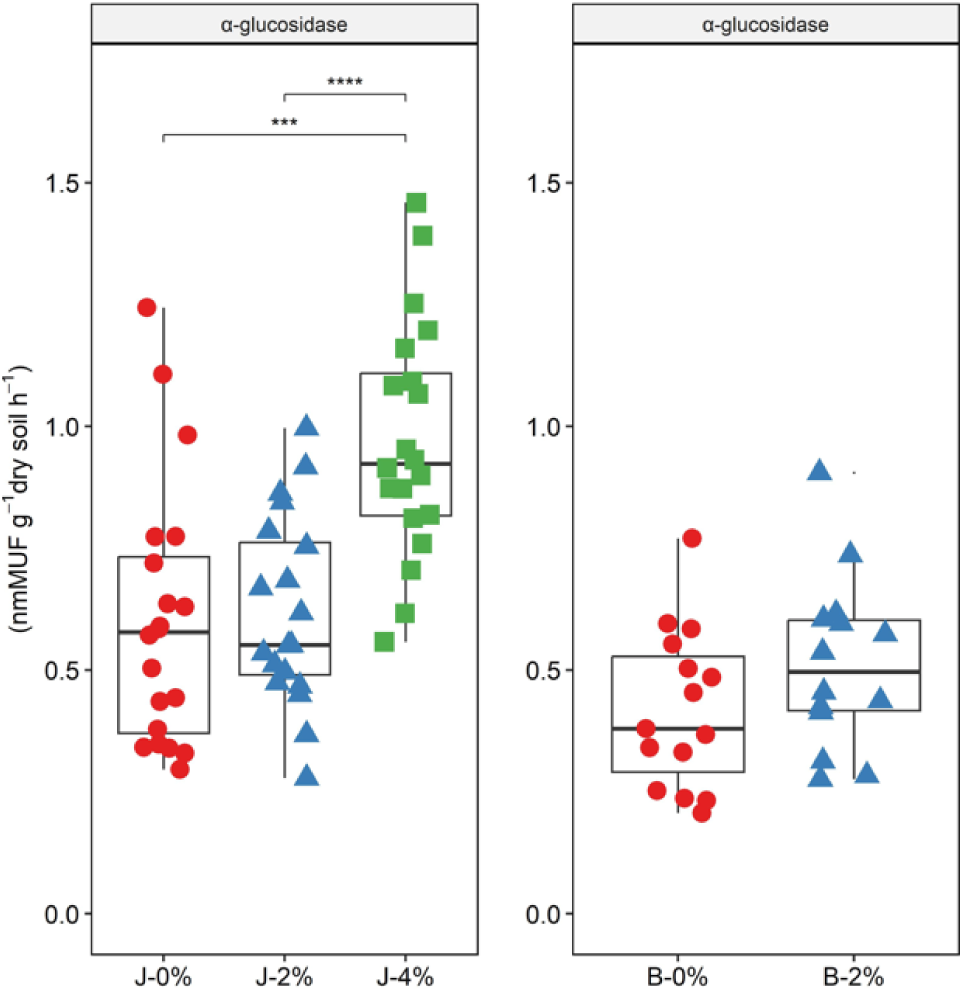
Extracellular α-glucosidase activity at 50 cm depth at the end of the experiment (August 2023). The unit nmMUF is an abbreviation for nanomoles of 4-methylumbelliferyl per gram of dry soil per hour. For treatment abbreviations see figure 3.

## DISCUSSION

### Water availability and yields

The incorporation of fine biochar particles into the coarse sandy soil increased subsoil water retention as hypothesized (Figure 2CD), whereas the addition of biochar pellets (J-2%-PEL) did not. In both growing seasons, the water content at field capacity in the biochar-amended subsoil (25-75 cm depth) generally increased with the addition of fine biochar, with the greatest increase of 25 mm in the treatment with the highest biochar dose (J-4%) compared to the non-irrigated control (J-0%) (Figure 2CD). Our findings of increased water retention following the addition of fine biochar agree with those of Bruun et al. (2014, 2021, 2023), although the effects observed in our present study are slightly lower. One reason could be less efficient mixing of the soil and biochar in the present study than in the other studies. Previously, we investigated the impact of different biochar particle sizes in the same coarse sandy soil under controlled laboratory conditions (Bruun et al. 2023). We documented substantial significant effects on water retention and hydraulic conductivity for particles with an average median diameter of 44 µm or less, while no or only limited effects were observed for larger biochar particles (> 81 µm) and biochar pellets. The increase in water retention and hydraulic conductivity was attributed to the particles converting drainable soil pores into smaller water-retaining pores. In our current study, this incorporation is also evidenced by the bulk density of the coarse sandy soil, which increased significantly with the addition of the small biochar particles, indicating that fine biochar particles settled into existing soil pores with little or no expansion of the soil particle matrix (Figure s-6).

However, in the first growing season (2022), despite the observed positive effect on soil water retention in the biochar-enriched layer (25-75 cm depth), barley yields were lower in the biochar treatments compared to the respective controls (Figure 3). Thus, for this year, our overall hypothesis that fine-grained biochar would improve the utilization of water and nutrients under drought conditions and thereby increase yield was not supported. This shows that the increased water retention in the biochar amended subsoil was not the primary factor determining crop yield in that year. However, in the following dry growing season of 2023, after intensive irrigation during the winter of 2022-23, the yield effects in the biochar treatments were either positive in the Billund soil or neutral in the Jyndevad soil. This suggests that the initial negative effects had been reduced or eliminated and that the yield effects in the second year were probably related to both water and nutrient availability.

In the second growing season, lack of water had notably negative impacts on both plant height and yields in all the non-irrigated treatments compared to the irrigated treatments. This reflects a more severe and earlier drought in 2023, which was already evident during the stem elongation growth phase, as also supported by the water balance data (Figure 2AB). However, the higher water content in the biochar-amended Billund soil (B-2%) was probably the main reason for the 54% increase in grain yield compared to B-0%, although this was still a very low yield. Under practical conditions, an additional 25 mm of plant-available water can be significant, allowing for improved water and nutrient utilization. However, this increase is relatively small compared to the amount of water retained at tensions below the wilting point in the deeper unamended layer of the soil columns (105-140 cm depth), estimated to be at least 78 mm. The barley roots observed at the bottom of all columns at the end of our experiment in August 2023 and the very low water contents measured near the lower boundary in July 2023 show that the plants in the non-irrigated treatments have had time to adapt to the drought and extract water down to the artificial lower boundary at about 140 cm depth. This could explain why we did not see clear effects of finely ground biochar on water extraction from the 0-105 cm layer (Figure 2 A, B). In the field, roots typically grow only to a maximum depth of 50 cm in soil with coarse sandy subsoil (e.g. Madsen and Platou, 1983). The deep roots observed in our experiment were probably a consequence of lower mechanical resistance to root growth due to the (unintended) lower subsoil bulk density (Figure s-6) in the soil columns (1.38 g cm^-3^ in J-0%) compared to the bulk density in the field (approximately 1.50 g cm^-3^, Hansen et al. 1986). Additionally, some preferential root growth occurred along the sides of the columns despite efforts to prevent this. As a result, when the upper part of the soil column began to dry out during the induced droughts, the higher water content in the biochar-enriched layer compared to the respective controls was not as critically important to the plants as it could have been under field conditions. Therefore, the experimental setup did not fully reflect the conditions that the roots would be exposed to under field conditions.

### Nitrogen availability after biochar application

Biochar influences N availability by affecting soil N turnover and transport, both directly and indirectly (Bruun et al. 2012; Liang et al. 2014; Xingren et al. 2020). Although biochar contains only a small proportion of labile carbon, its addition to soil, especially at high doses, can stimulate microbial activity, as also indicated by the higher extracellular α-glucosidase activity in the treatments with biochar (J-4%) in the current study (Figure 9). However, due to the low N content of biochar, soil microbes may experience N deficiency, leading them to scavenge additional N from the soil (Bruun et al. 2012; Hansen et al. 2015). In the first season of our experiment, the reduced biomass yields and plant heights observed in the treatments with biochar coincided with significantly lower total N uptake and concentration in both grains and straw (Figure 4). This occurred despite the application of NPK fertilizer (equivalent to 200 kg N/ha) to all treatments before both growing seasons. A similar effect was observed in the two irrigated treatments (J-0%-IR and J-2%-IR), where the irrigated control produced 10% more grain than the biochar-amended irrigated treatment. This suggests that N immobilization in the soil may have played a role in the observed yield reductions (Figure 3), supporting our second hypothesis that the application of biochar may inhibit plant growth in the initial months due to N deficiency.

In the second season, the non-irrigated biochar treatments also had a tendency to lower grain N concentrations (only J-2%-PEL was significant) in plant biomass compared to the control (J-0%). However, the potential yield reductions were likely outweighed by the slightly higher subsoil water content compared to the respective controls. Over time, as the labile carbon fraction of the biochar is mineralized, the risk of nitrogen immobilization may diminish.

### Phosphorus availability after biochar application

During the first growing season, phosphate (P) concentrations in the grains were similar between the irrigated and non-irrigated treatments (Figure 5AB). However, during the second growing season, low grain P concentrations in the non-irrigated treatments indicated reduced P uptake by roots due to more severe drought conditions in both the topsoil and subsoil.

Interestingly, biochar did not significantly affect grain P concentration in either the irrigated or non-irrigated treatments, despite the substantial P contribution from biochar, which e.g. in J-4% added about 88 mg total P/kg soil to the subsoil. This indicates that biochar P did not substantially contribute to plant P supply (Table s-1). One reason for this could be the reduced root density associated with biochar application. However, low plant P availability is also indicated by the low levels of Olsen P measured in the subsoil in all treatments, with no consistent trends over the two seasons (Figure 8B). Biochar derived from lignocellulosic feedstocks has been proposed as an alternative P fertilizer (Robinson et al. 2018), but as the dominant forms of P in these materials are Ca-bound phosphates, they will have the greatest fertilizing effect in acidic soils (Sun et al. 2018), which was not the condition in our experiment. Furthermore, it has often been shown that P availability from biochar is greatest when produced at moderate pyrolysis temperatures (i.e. below 600 °C) (Glaser & Lehr 2019). Considering the tested application of high doses of biochar to the subsoil, the observed low P solubility may also be an advantage, as the risk of P leaching is likely to be minimal (Wang et al. 2015).

### Other effects on plant growth

Despite roots at the bottom of all columns, the root density (estimated as the root coverage at selected depths) appeared to be negatively affected by biochar (Figure 6). The root coverage decreased significantly with increasing biochar concentrations at all depths in the Jyndevad soil compared to the control, although this effect was less clear in the Billund soil. The exact reasons for these reductions in root density remain unclear. Soil pH levels were not exceptionally high, even in the high dose biochar treatment (J-4%) with 7.3 (Figure 7B) and are thus unlikely to have reduced root growth. However, high potassium levels in the biochar treatments, combined with the low soil water content, may have caused plant salinity stress and inhibited plant and root growth (Neumann, 1995). In the first season, exchangeable K levels in the J-4% biochar treatment were up to 14 times higher than in the control soil, with the control showing relatively low concentrations (ranging from 15 to 29 mg kg^-1^ in the subsoil over the two years). While potassium (K) is essential for processes such as osmotic regulation and stomatal function (Wang et al. 2013), excess K can cause osmotic stress and nutrient imbalances by suppressing the uptake of calcium (Ca) and magnesium (Mg) (Butnan et al. 2015; Buss et al. 2016). In a study by Ruan et al. (2024) investigating the effect of biochar on maize root growth, a reduction in lateral roots was observed. The researchers attributed this to elevated K levels following biochar application, which increased K concentrations in the root sap while reducing Ca and Na levels in two maize genotypes compared to the control. They suggested that the higher K levels increased the apoplastic pH, thereby inhibiting lateral root development. In the present study, higher K concentrations in the biochar-amended soils were associated with increased K levels in the straw (Figure 5C, Table s-3), but not in the grains.

Conversely, calcium concentrations in the grain were consistently reduced in the biochar treatments (Figure 5F), while magnesium concentrations remained unaffected, probably because most nutrient uptake occurred from the fertilized topsoil. Despite the high potassium concentrations, electrical conductivity (EC), ranging from 0.03 to 0.08 mS/cm, indicated non-saline conditions in the sandy soil (Hardie & Doyle, 2012) and was comparable to EC values reported previously (Bruun et al. 2021). Following intensive irrigation in winter 2022–2023—with a percolation equivalent to three years’ average net precipitation—both EC levels and exchangeable K decreased significantly, although the latter was still higher in the biochar treatments compared to the control. Thus, it is very likely that these levels would continue to decrease in subsequent years. Nevertheless, the initial K content of biochar should be carefully considered when selecting application rates and feedstocks, e.g. wood-based biochars typically contain much less K than those derived from straw.

In addition to the effects mentioned above, typical negative effects on root growth in coarse sandy soil include mechanical resistance to root growth. The lower bulk density in the Jyndevad control soil (J-0%, 1.38 g cm^-3^) compared to the biochar treatments (J-X%, 1.39-1.43 g cm^-3^) likely resulted in lower mechanical resistance in the control soil which may have contributed to the better root growth observed here. However, these results may not fully represent field conditions, because the bulk density is typically higher than in the current study, as mentioned above.

## CONCLUSION

In this study, we evaluated the impact of high, single applications of biochar (up to 300 Mg/ha) to coarse sandy subsoil on spring barley. Our results showed that finely ground biochar increased the available water capacity (AWC), supporting results from previous laboratory experiments. However, in the first growing season of our study (2022), we observed negative effects on plant height, biomass yield, and grain nitrogen concentration in the biochar treatments. In the dry 2023 growing season, after intensive irrigation during the winter between 2022 and 2023, the yield effects of the biochar applications were either positive or neutral. This suggests that initial negative effects had been reduced or eliminated and that yield effects in the second year were related to both water and nutrient availability. However, root density was negatively affected by biochar, especially in one of the soils tested. The exact cause remains unclear, but increased levels of potassium in the subsoil originating from the high doses of biochar may have created unfavourable conditions.

The effects of biochar on plant yields and development observed in this study should be interpreted as short-term results of its application to coarse sandy subsoil, because biochar contains labile fractions and salts that can initially alter soil properties but may be leached or mineralized in the first months after application. The specific reasons for the negative effects of high subsoil applications of biochar on plant growth should be further investigated in longer-term field experiments but may be mitigated by choosing the right timing of application, type of biochar and additional supply of plant nutrients. As subsoil incorporation of biochar would require a large-scale operation, specialized machinery will need to be developed to reduce costs and ensure thorough mixing of fine biochar into sandy subsoil.

## ACKNOWLEDGEMENTS

We are grateful to Stiesdal A/S for providing the biochar, and to Giulia Ravenni for assistance with various analyses and for facilitating the grinding of the biochar pellets. We also thank Simon Svane and Andrea Parker for technical assistance.

## REFERENCES

Andersen MN (1985) Planternes tørkeresistens, rodudvikling og vandforråd på sandjord. Tidsskrift for Planteavls Specialserie, Aarhus University, Aarhus.

Brtnicky M, Datta R, Holatko J, Bielska L, Gusiatin ZM, Kucerik J, … Pecina V (2021) A critical review of the possible adverse effects of biochar in the soil environment. Science of the Total Environment 796, 148756.

Bruun EW, Ambus P, Egsgaard H, Hauggaard-Nielsen H (2012) Effects of slow and fast pyrolysis biochar on soil C and N turnover dynamics. Soil Biology and Biochemistry 46, 73–79.

Bruun EW, Petersen CT, Hansen E, Holm JK, Hauggaard-Nielsen H (2014) Biochar amendment to coarse sandy subsoil improves root growth and increases water retention. Soil Use and Management 30(1), 109–118.

Bruun EW, Müller-Stöver D, Pedersen BN, Hansen LV, Petersen CT (2021) Ash and biochar amendment of coarse sandy soil for growing crops under drought conditions. Soil Use and Management 38(2), 1280–1292.

Bruun EW, Ravenni G, Müller-Stöver D, Petersen CT (2023) Small biochar particles added to coarse sandy subsoil greatly increase water retention and affect hydraulic conductivity. European Journal of Soil Science 74(6), e13442.

Buss W, Assavavittayanon K, Shepherd JG, Heal KV, Sohi S (2018) Biochar phosphorus release is limited by high pH and excess calcium. Journal of Environmental Quality 47(5), 1298–1303.

EBC (2012–2023) European Biochar Certificate – Guidelines for a Sustainable Production of Biochar. Carbon Standards International (CSI), Frick, Switzerland. Version 10.3 from 5th Apr 2022. Available at: http://european-biochar.org

Gill JS, Clark GJ, Sale PW, Peries RR, Tang C (2012) Deep placement of organic amendments in dense sodic subsoil increases summer fallow efficiency and the use of deep soil water by crops. Plant and Soil 359, 57–69.

Glaser B, Lehr VI (2019) Biochar effects on phosphorus availability in agricultural soils: A meta-analysis. Scientific Reports 9, 9338.

Hansen V, Hauggaard-Nielsen H, Petersen CT, Mikkelsen TN, Müller-Stöver D (2016) Effects of gasification biochar on plant-available water capacity and plant growth in two contrasting soil types. Soil and Tillage Research 161(5), 1–9. 10.1016/j.still.2016.03.002

Hansen S, Storm B, Jensen HE (1986) Spatial variability of soil physical properties theoretical and experimental analyses – Soil sampling, experimental analyses and basic statistics of soil physical properties. Research Report nr. 1201, The Royal Veterinary and Agricultural University, Copenhagen.

Hardie M, Doyle R (2012) Measuring soil salinity. In: Shabala S (Ed.) Plant Salt Tolerance: Methods and Protocols, 415–425.

Henriksen HJ, Sonnenborg A (2003) Ferskvandets kredsløb. NOVA Temarapport, GEUS.

Iturbe-Espinoza M, et al. (2025) In review.

Joseph S, Cowie AL, Van Zwieten L, Bolan N, Budai A et. Al. (2021) How biochar works, and when it doesn’t: A review of mechanisms controlling soil and plant responses to biochar. GCB Bioenergy 13(11), 1731–1764.

Larsen V, Øvig JK (1979) Halm til jordforbedring af sandjord. Meddelelse. Hedeselskabets Forsøgsvirksomhed (Denmark) no. 8.

Lehmann J, Abiven S, Kleber M, Pan G, Singh BP, Sohi SP, Zimmerman AR, Lehmann J, Joseph S (2015) Persistence of biochar in soil. In: Lehmann J, Joseph S (Eds.) Biochar for Environmental Management: Science, Technology and Implementation (Vol. 2), 233–280. Routledge.

Madsen HB, Platou SW (1983) Land use planning in Denmark. Nordic Hydrology 14, 267–276.

Munkholm LJ, Schjønning P, Sørensen H (2003) Jordpakning og mekanisk løsning på grovsandet jord. Grøn Viden – Markbrug 271.

Munkholm LJ, Schjønning P, Jørgensen MH, Thorup-Kristensen K (2005) Mitigation of subsoil recompaction by light traffic and on-land ploughing: II. Root and yield response. Soil and Tillage Research 80(1–2), 159–170.

Munkholm LJ, Hansen EM, Olesen JE (2008) The effect of tillage intensity on soil structure and winter wheat root/shoot growth. Soil Use and Management 24, 392–400.

Møberg JP, Petersen L, Rasmussen K (1988) Constituents of some widely distributed soils in Denmark. Geoderma 42, 295–316.

Petersen CT, Hansen E, Larsen HH, Hansen LV, Ahrenfeldt J, Hauggaard-Nielsen H (2016) Pore-size distribution and compressibility of coarse sandy subsoil with added biochar. European Journal of Soil Science 67(6), 726–736.

Robinson JS, Baumann K, Hu Y, Hagemann P, Kebelmann L, Leinweber P (2018) Phosphorus transformations in plant-based and bio-waste materials induced by pyrolysis. Ambio 47, 73–82.

Schmidt HP, Wilson K (2012) 55 uses of biochar. Ithaka Journal 1, 286–289.

Skatteministeriet (2024) Grøn skattereform – endelig afrapportering. Available at: https://www.skm.dk/aktuelt/publikationer/rapporter/groen-skattereform-endelig-afrapportering

Sun K, Qiu M, Han L, Jin J, Wang Z, Pan Z, Xing B (2018) Speciation of phosphorus in plant- and manure-derived biochars and its dissolution under various aqueous conditions. Science of the Total Environment 634, 1300–1307.

Thengane SK, Bandyopadhyay S (2020) Biochar mines: Panacea to climate change and energy crisis? Clean Technologies and Environmental Policy 22(1), 5–10.

Wang Y, Lin Y, Chiu PC, Imhoff PT, Guo M (2015) Phosphorus release behaviors of poultry litter biochar as a soil amendment. Science of the Total Environment 512, 454–463.

Woolf D, Lehmann J, Cowie A, Cayuela ML, Whitman T, Sohi S (2018) Biochar for climate change mitigation. In: Lal R, Stewart BA (Eds.) Soil and Climate, 219–248.

Øvig JK (1979) Forsøg med sphagnumgødning. Beretning nr. 20, Det danske Hedeselskab, Viborg.

