## Supplementary material for "High doses of fine biochar in sandy subsoils increase water retention, but also cause first-year yield depressions for drought-stressed barley"

**SUPPLEMENTARY MATERIALS**

*Analyses of enzymatic activity*

For analyses of enzymatic activity soils were sieved (2 mm mesh) and 5 g of each soil was mixed with 50 mL of MQ water in 250 mL centrifuge tubes. The tubes were subjected to cold ultrasonic treatment in an ice bath for 5 min, followed by shaking at 1000 rpm for 5 min in a Geno/Grinder 2000 (SPEX CertiPrep, Metuchen, NJ, USA). After centrifuging at 750 rpm for 10 minutes at 4°C (J2-21M, rotor JA-14, Beckman USA), supernatants were collected and stored at 4°C until enzymatic assays. The assays were conducted in 96-well microtiter plates using 200 µL of soil supernatant, 10 µL MOPS buffer (pH 7.4), and 40 µL of 50 µM 4-Methylumbelliferyl-α-D-glucoside (α-glucosidase, EC 3.2.1.20). Calibration curves were created with different concentrations of 4-methylumbelliferone, 7-hydroxy-4-methylcoumarin sodium salt (62.4 µM, 31.2 µM, 15.6 µM, and 0 µM) for each soil sample. Fluorescence (excitation at 360 nm with a bandwidth of 20 nm, emission at 450 nm with a bandwidth of 30 nm) was measured every 10 min for 10 cycles (CLARIOstar Plus, Germany), and enzyme activity was calculated based on the linear regression slope of the standard curve and expressed in nmMUF $g^{-1}$ dry soil $h^{-1}$.

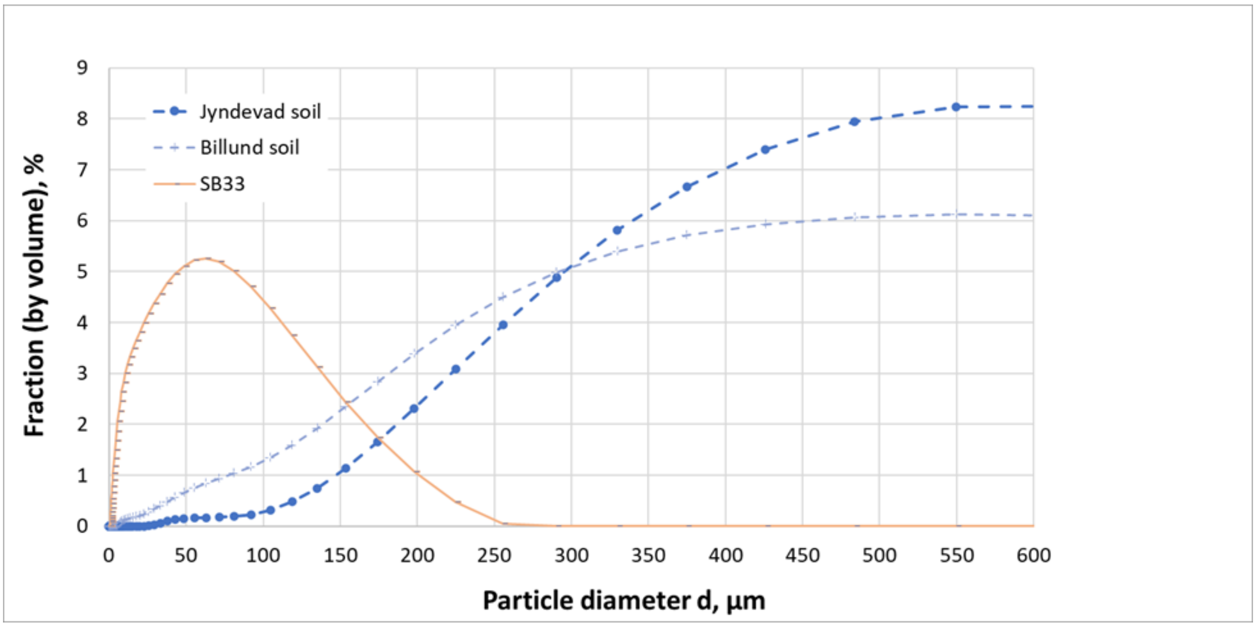


*Figure s-1. Particle size distribution curves for the fine biochar used in the experiment and for the coarse sandy (Jyndevad soil).*

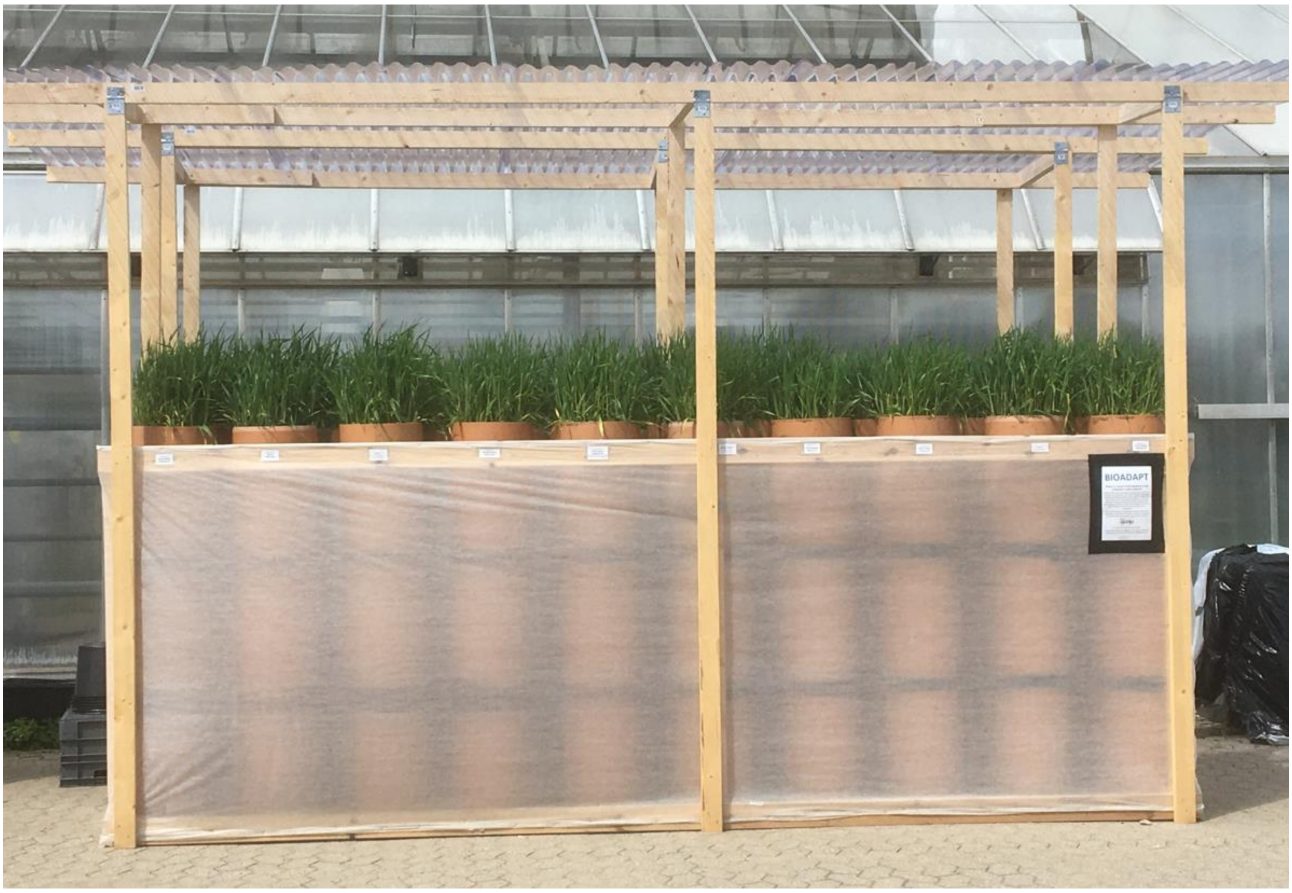


*Figure s-2. Experimental setup with rain shelter roof. The outmost row with the randomized eight treatments (Block D). Photo of June 8, 2022.*

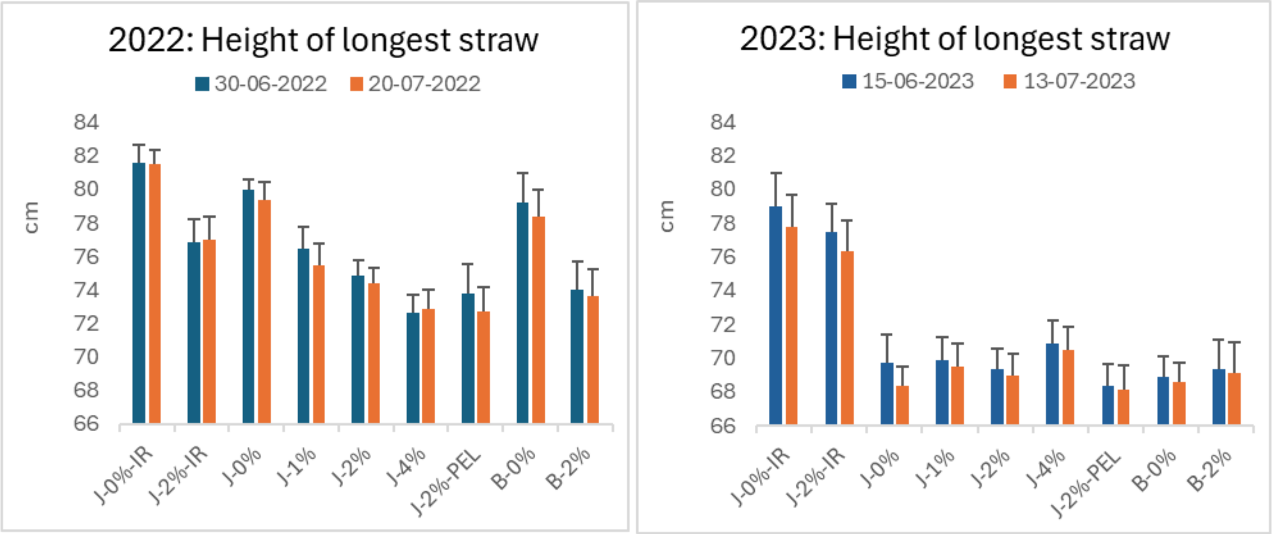


*Figure s-3. Average length of longest shoot measured at different dates in 2022 (A) and in 2023 (B). Mean ± standard error.*

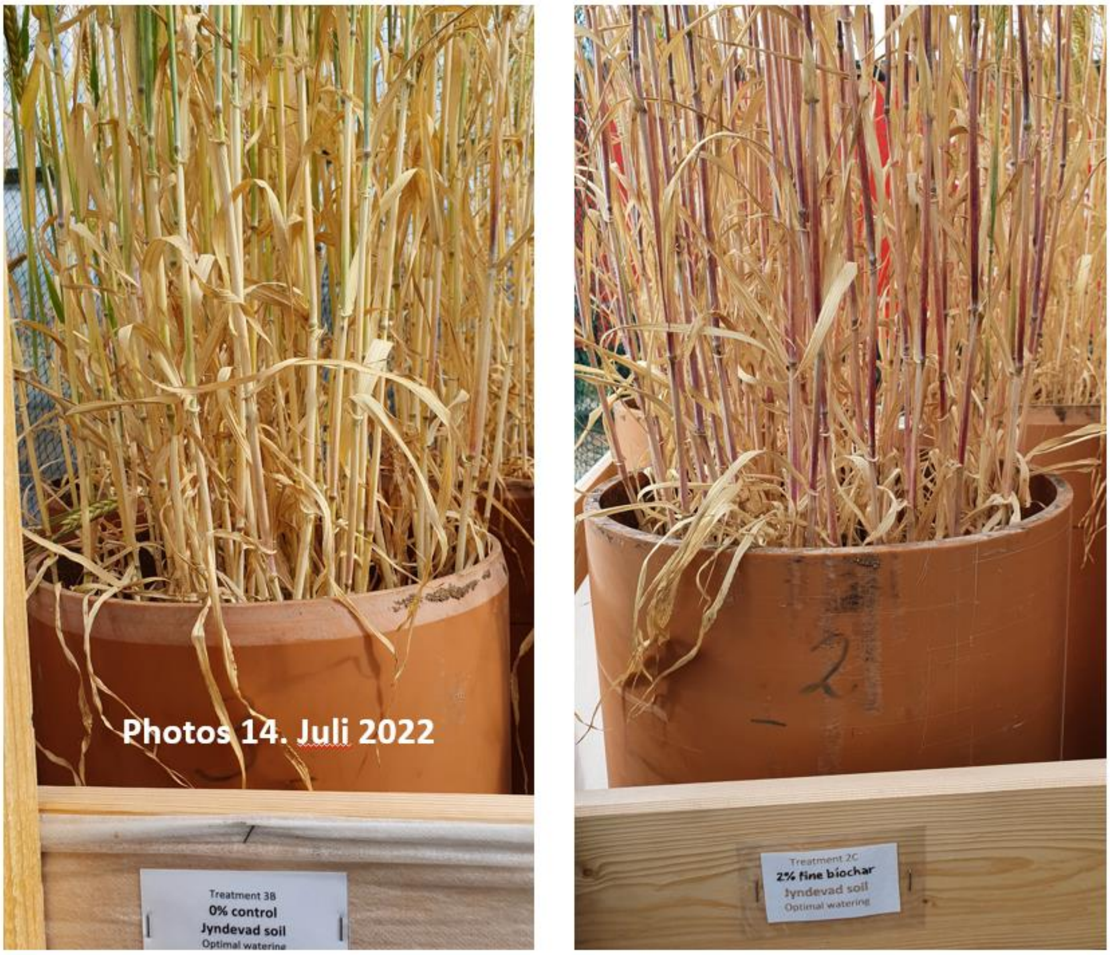


*Figure s-4. Reddish color on stems in treatments with biochar. Photo from July 14, 2022.*

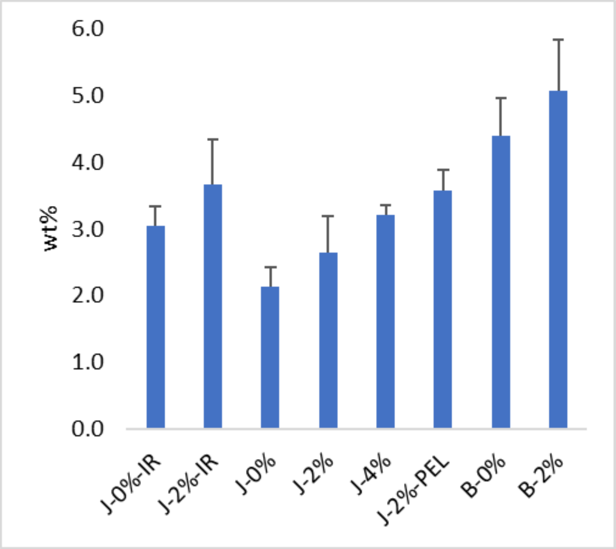


*Figure s-5. Water content measured July 13-2023 in all columns at appr. 130-140 cm's depth (soil samples were obtained 15 cm above ground in each column).*

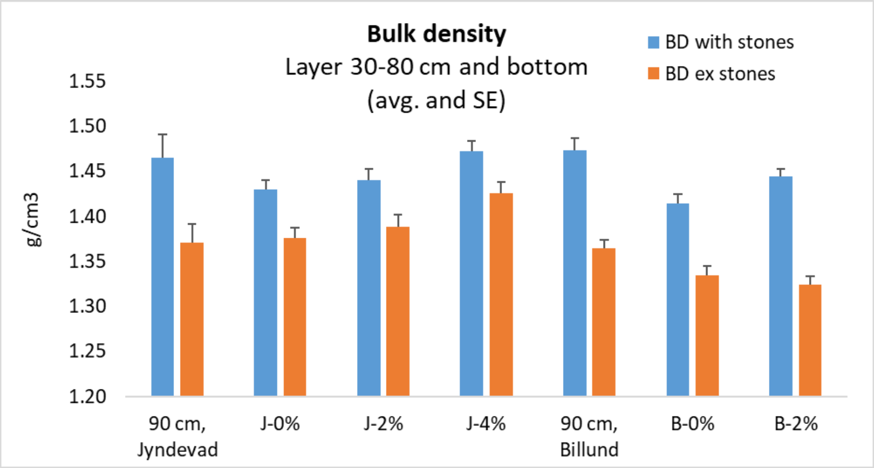


*Figure s-6. Bulk density for the investigated samples (n = 3) of subsoil with and without biochar.*

Table s-1. Elemental composition and nutrients added with fine grained biochar at 2% and 4%.

| Nutrients added with the biochar | µg /g | | mg/kg subsoil | |
|---|---|---|---|---|
| | Avg. | SE | 2% biochar | 4% biochar |
| K | 25853 | 500 | 517 | 1034 |
| Ca | 8085 | 121 | 162 | 323 |
| P | 2200 | 47 | 44 | 88 |
| Mg | 1835 | 13 | 37 | 73 |
| S | 1936 | 43 | 39 | 77 |
| Al | 475 | 14 | 10 | 19 |
| Na | 467 | 2 | 9 | 19 |
| Fe | 353 | 18 | 7 | 14 |
| Mn | 120 | 1 | 2 | 5 |
| Zn | 63 | 2 | 1 | 3 |
| Cu | 6.5 | 1.3 | 0.13 | 0.26 |
| Cr | 4.7 | 0.7 | 0.09 | 0.19 |
| Cd | 0.72 | | 0.01 | 0.03 |
| B | nd | - | - | - |

Table s-2. PAH content analysed by Eurofins (2022-04-11, Eurofins, Bobritzsch-Hilbersdorf).

eurofins Umwelt



| | | | | Limit values | | | | | | Description | | straw biochar 30-03-2022 | |
|---|---|---|---|---|---|---|---|---|---|---|---|---|---|
| | | | | | | | | | | Date and time of sample taking | | 2022-03-30 | |
| | | | | | | | | | | Sample number | | 122049298 | |
| **Parameter** | **Lab** | **Accr.** | **Method** | EBC-Feed | EBC-Agro Organic | EBC-Agro | EBC-Urban | EBC-Con-sumer Materials | EBC-Basic Materials | **LOQ** | **Unit** | **ar** | **db** |
| Total 8 EFSA-EPA excl. LOQ | FR | RE000 FY | DIN EN 16181:2019-08 | 1 | 1 | 1 | 1 | 1 | 4 | | mg/kg | - | 0.8 |
| Total 16 EPA-PAH excl. LOQ | FR | RE000 FY | DIN EN 16181:2019-08 | | 4[1)] | 6[1)] | | | | | mg/kg | - | 3.7 |
| Benzo(e)pyrene | FR | RE000 FY | DIN EN 16181:2019-08 | < 1 | < 1 | < 1 | < 1 | < 1 | < 1 | 0.1 | mg/kg | - | 0.1 |
| Benzo-(j)-fluoranthen | FR | RE000 FY | DIN EN 16181:2019-08 | < 1 | < 1 | < 1 | < 1 | < 1 | < 1 | 0.1 | mg/kg | - | < 0.1 |

### Explanations

LOQ - Limit of quantification
ar - as received
db - dry basis
Lab - Abbreviation of the performing laboratory
Accr. - Abbreviation of the accreditation of the performing laboratory

The parameters identified by FR have been performed by the laboratory Eurofins Umwelt Ost GmbH (Lindenstraße 11, Gewerbegebiet Freiberg Ost, Bobritzsch-Hilbersdorf). The accreditation code RE000FY identifies the parameters accredited according to DIN EN ISO/IEC 17025:2018 DAkkS D-PL-14081-01-00 .

Table s-3. Nutrients in straw. Concentrations of selected mineral nutrients in barley straw after harvest (10-08-2022 and 13-07-2023). Numbers in parenthesis are standard errors.

| **STRAW** | 2022 | 2023 | 2022 | 2023 | 2022 | 2023 | 2022 | 2023 | 2022 | 2023 | 2022 | 2023 | 2022 | 2023 |
|---|---|---|---|---|---|---|---|---|---|---|---|---|---|---|
| | P µg /g | | Ca µg /g | | K µg /g | | Mg µg /g | | Mn µg /g | | Cu µg /g | | Zn µg /g | |
| J-0%-IR | 233 (15) | 240 (15) | 3150 (183) | 6089 (289) | 12693 (943) | 12664 (435) | 673 (22) | 1641 (70) | 25 (2) | 7.5 (0.5) | 2.0 (0.1) | 2.1 (0.1) | 27 (4) | 17 (1) |
| J-2%-IR | 269 (17) | 259 (26) | 3402 (119) | 5395 (447) | 17404 (217) | 16686 (202) | 743 (28) | 1469 (71) | 31 (4) | 7 (0.3) | 1.8 (0.1) | 2.0 (0.1) | 31 (5) | 18 (1) |
| J-0% | 245 (6) | 241 (7) | 3638 (222) | 6749 (277) | 11885 (699) | 11548 (572) | 749 (48) | 1726 (77) | 34 (3) | 14.6 (0.4) | 2.2 (0.1) | 2.5 (0.1) | 23 (2) | 33 (1) |
| J-2% | 263 (44) | 204 (6) | 3238 (180) | 5965 (297) | 15796 (899) | 14571 (685) | 733 (39) | 1564 (70) | 43 (3) | 14.5 (0.4) | 2.2 (0.3) | 2.4 (0) | 33 (5) | 33 (1) |
| J-4% | 275 (53) | 252 (33) | 2774 (325) | 6884 (344) | 19541 (497) | 15997 (768) | 606 (58) | 1784 (97) | 37 (1) | 15 (0.8) | 2.0 (0.1) | 2.6 (0.1) | 32 (6) | 36 (2) |
| J-2%-PEL | 231 (28) | 193 (6) | 2583 (237) | 6426 (486) | 17520 (508) | 15039 (297) | 660 (32) | 1601 (88) | 48 (5) | 16.5 (0.7) | 1.8 (0.2) | 2.4 (0) | 35 (5) | 32 (1) |
| B-0% | 246 (28) | 221 (10) | 5396 (668) | 10379 (1073 | 11741 (1003 | 11057 (663) | 824 (55) | 1752 (143) | 24 (2) | 15.7 (1.2) | 2.4 (0) | 2.5 (0.1) | 14 (1) | 19 (0) |
| B-2% | 259 (11) | 228 (26) | 3759 (138) | 9124 (320) | 21064 (933) | 15360 (645) | 711 (19) | 1562 (62) | 25 (1) | 16.5 (0.8) | 2.0 (0.1) | 2.2 (0.1) | 14 (2) | 20 (1) |
| **STRAW** | 2022 | 2023 | 2022 | 2023 | 2022 | 2023 | 2022 | 2023 | 2022 | 2023 | 2022 | 2023 | 2023 | 2023 |
| | Fe µg /g | | Na µg /g | | S µg /g | | Sr µg /g | | Al µg /g | | V µg /g | Cd µg /g | Cr µg /g | B µg /g |
| J-0%-IR | 79 (5) | 93 (12) | 893 (48) | 1579 (65) | 1408 (85) | 2127 (124) | 6.5 (0.2) | 10.5 (0.5) | 88 (5) | 78 (10) | 0.43 (0.01) | 0.77 (0.02) | 0.58 (0.15) | 5.9 (0.2) |
| J-2%-IR | 87 (10) | 97 (10) | 667 (87) | 1318 (77) | 1367 (107) | 2247 (70) | 8.4 (0.4) | 9.9 (0.6) | 85 (10) | 80 (9) | 0.37 (0.02) | 0.71 (0.02) | 0.76 (0.12) | 6.1 (0.5) |
| J-0% | 104 (14) | 120 (4) | 1226 (55) | 1118 (45) | 1240 (65) | 1295 (32) | 6.7 (0.3) | 10.1 (0.5) | 115 (20) | 103 (5) | 0.34 (0.02) | 0.8 (0.02) | 1.13 (0.12) | 4.5 (0.2) |
| J-2% | 102 (23) | 98 (8) | 736 (18) | 1041 (61) | 1274 (102) | 1304 (21) | 6.8 (0.2) | 9.7 (0.6) | 97 (23) | 83 (9) | 0.42 (0.02) | 0.75 (0.02) | 0.89 (0.12) | 4.3 (0.1) |
| J-4% | 80 (16) | 113 (9) | 875 (80) | 1218 (86) | 1199 (108) | 1498 (43) | 6.4 (0.7) | 11.0 (0.7) | 80 (14) | 96 (9) | 0.40 (0.02) | 0.84 (0.01) | 0.74 (0.1) | 4.8 (0.2) |
| J-2%-PEL | 80 (16) | 116 (10) | 697 (25) | 1021 (37) | 1162 (77) | 1341 (25) | 5.6 (0.5) | 10.1 (0.8) | 83 (14) | 98 (9) | 0.37 (0.02) | 0.79 (0.03) | 0.97 (0.11) | 4.4 (0.2) |
| B-0% | 100 (10) | 92 (12) | 914 (74) | 1082 (54) | 1548 (65) | 1308 (69) | 11.4 (1.2) | 21.0 (2.3) | 111 (11) | 77 (9) | 0.39 (0.04) | 0.92 (0.06) | 0.80 (0.21) | 4.6 (0.2) |
| B-2% | 72 (13) | 91 (7) | 612 (81) | 1017 (82) | 1490 (81) | 1361 (52) | 10.1 (0.2) | 18.6 (0.9) | 72 (12) | 75 (6) | 0.44 (0.02) | 0.89 (0.02) | 0.64 (0.02) | 5.0 (0.2) |

Table s-4. Nutrients in grains. Concentrations of selected mineral nutrients in barley grains after harvest (10-08-2022 and 13-07-2023). Numbers in parenthesis are standard errors.

| **GRAIN** | 2022 | 2023 | 2022 | 2023 | 2022 | 2023 | 2022 | 2023 | 2022 | 2023 | 2022 | 2023 | 2022 | 2023 |
|---|---|---|---|---|---|---|---|---|---|---|---|---|---|---|
| | P (µg/g) | | Ca (µg/g) | | K (µg/g) | | Mg (µg/g) | | Mn (µg/g) | | Cu (µg/g) | | Zn (µg/g) | |
| J-0%-IR | 2342 (73) | 1823 (39) | 365 (15) | 334 (9) | 4094 (107) | 4391 (140) | 1043 (26) | 951 (23) | 16 (1) | 6.9 (0.5) | 4.6 (0.4) | 2.5 (0.1) | 34 (1) | 27 (1) |
| J-2%-IR | 2434 (94) | 1960 (76) | 360 (6) | 315 (10) | 4086 (113) | 4651 (196) | 1028 (23) | 971 (13) | 15 (0) | 5.8 (0.9) | 5.0 (0) | 2.5 (0.1) | 33 (1) | 26 (0) |
| J-0% | 2679 (77) | 1188 (31) | 402 (13) | 442 (11) | 4695 (90) | 4728 (111) | 1082 (16) | 729 (9) | 19 (0) | 10.1 (0.3) | 5.0 (0.1) | 2.4 (0) | 43 (1) | 41 (1) |
| J-2% | 2293 (63) | 1097 (25) | 334 (8) | 388 (9) | 4155 (164) | 4614 (78) | 1031 (23) | 737 (14) | 19 (1) | 10.5 (0.3) | 5.1 (0.1) | 2.5 (0.1) | 42 (2) | 40 (2) |
| J-4% | 2583 (88) | 1142 (37) | 316 (7) | 336 (10) | 4357 (128) | 4569 (122) | 1053 (17) | 679 (24) | 18 (1) | 9.9 (0.3) | 4.9 (0.3) | 2.2 (0) | 40 (1) | 35 (1) |
| J-2%-PEL | 2359 (42) | 981 (57) | 334 (13) | 354 (9) | 4286 (260) | 4691 (120) | 1065 (24) | 694 (32) | 20 (1) | 10.0 (0.2) | 5.5 (0.5) | 2.2 (0.1) | 43 (3) | 35 (2) |
| B-0% | 2088 (72) | 1049 (33) | 446 (51) | 460 (14) | 3946 (69) | 4474 (136) | 984 (26) | 798 (18) | 15 (0) | 10.3 (0.1) | 4.5 (0.2) | 2.3 (0.2) | 32 (3) | 32 (0) |
| B-2% | 2307 (21) | 1169 (75) | 334 (9) | 379 (7) | 3701 (55) | 4492 (95) | 982 (15) | 746 (5) | 16 (2) | 10.6 (0.2) | 4.8 (0.4) | 2.6 (0.2) | 28 (2) | 34 (1) |
| **GRAIN** | 2022 | 2023 | 2022 | 2023 | 2022 | 2023 | 2022 | 2023 | 2022 | 2023 | 2022 | 2023 | 2023 | 2023 |
| | Fe (µg/g) | | Na (µg/g) | | S (µg/g) | | Sr (µg/g) | | Al (µg/g) | | V (µg/g) | Cd (µg/g) | Cr (µg/g) | B (µg/g) |
| J-0%-IR | 48 (3) | 38 (4) | 114 (11) | 175 (15) | 912 (22) | 1102 (8) | 1.01 (0.07) | 0.71 (0.03) | 61 (10) | 86 (2) | 0.37 (0.01) | 0.40 (0.02) | 0.90 (0.34) | 1.1 (0.1) |
| J-2%-IR | 46 (3) | 29 (1) | 74 (6) | 127 (8) | 834 (14) | 1063 (17) | 1.13 (0.04) | 0.72 (0.04) | 69 (1) | 87 (5) | 0.41 (0.04) | 0.39 (0.02) | 0.32 (0.17) | 0.9 (0.1) |
| J-0% | 57 (3) | 55 (2) | 114 (8) | 97 (3) | 1083 (21) | 1112 (29) | 0.92 (0.03) | 0.82 (0.02) | 62 (2) | 64 (4) | 0.42 (0.01) | 0.39 (0.01) | 0.80 (0.25) | 0.9 (0.1) |
| J-2% | 65 (6) | 57 (3) | 95 (9) | 96 (6) | 915 (31) | 1054 (13) | 0.78 (0.03) | 0.78 (0.03) | 61 (3) | 80 (5) | 0.4 (0.01) | 0.38 (0.01) | 0.80 (0.21) | 1.1 (0.1) |
| J-4% | 52 (3) | 46 (0) | 84 (9) | 96 (11) | 896 (23) | 1037 (29) | 0.79 (0.03) | 0.66 (0.03) | 61 (4) | 76 (5) | 0.38 (0.01) | 0.37 (0.01) | 0.49 (0.12) | 1.0 (0.1) |
| J-2%-PEL | 64 (6) | 51 (0) | 90 (11) | 107 (6) | 890 (37) | 955 (47) | 0.77 (0.03) | 0.72 (0.02) | 72 (8) | 80 (2) | 0.37 (0.01) | 0.40 (0.01) | 1.32 (0.28) | 1.0 (0.1) |
| B-0% | 48 (8) | 44 (2) | 92 (7) | 90 (6) | 1075 (83) | 1111 (11) | 1.11 (0.13) | 1.2 (0.05) | 51 (4) | 46 (4) | 0.35 (0.01) | 0.36 (0.02) | 0.60 (0.43) | 1.0 (0.1) |
| B-2% | 36 (5) | 50 (3) | 56 (1) | 71 (2) | 876 (23) | 1091 (31) | 1.02 (0.03) | 1.01 (0.04) | 71 (13) | 76 (4) | 0.41 (0.01) | 0.36 (0.01) | 1.08 (0.61) | 0.9 (0.1) |
